# A family of novel immune systems targets early infection of nucleus-forming jumbo phages

**DOI:** 10.1101/2022.09.17.508391

**Authors:** Yuping Li, Jingwen Guan, Surabhi Hareendranath, Emily Crawford, David A. Agard, Kira S. Makarova, Eugene V. Koonin, Joseph Bondy-Denomy

## Abstract

Jumbo bacteriophages of the ⌽KZ-like family are characterized by large genomes (>200 kb) and the remarkable ability to assemble a proteinaceous nucleus-like structure. The nucleus protects the phage genome from canonical DNA-targeting immune systems, such as CRISPR-Cas and restriction-modification. We hypothesized that the failure of common bacterial defenses creates selective pressure for immune systems that target the unique jumbo phage biology. Here, we identify the “<u>ju</u>mbo phage <u>k</u>iller” (Juk) immune system that is deployed by a clinical isolate of *Pseudomonas aeruginosa* to resist ⌽KZ. Juk immunity rescues the cell by preventing early phage transcription, DNA replication, and nucleus assembly. Phage infection is first sensed by JukA (formerly YaaW), which localizes rapidly to the site of phage infection at the cell pole, triggered by ejected phage factors. The effector protein JukB is recruited by JukA, which is required to enable immunity and the subsequent degradation of the phage DNA. JukA homologs are found in several bacterial phyla and are associated with numerous other putative effectors, many of which provided specific anti-⌽KZ activity when expressed in *P. aeruginosa*. Together, these data reveal a novel strategy for immunity whereby immune factors are recruited to the site of phage protein and DNA ejection to prevent phage progression and save the cell.

## Introduction

The viruses that infect bacteria have evolved numerous strategies to ensure faithful replication, assembly, and lysis of their host in an exquisitely-timed manner. Conversely, bacteria employ a suite of diverse defense pathways to block phage injection, replication, or maturation. Anti-phage defenses can prevent adsorption/DNA ejection, directly act on the phage nucleic acid, or sense synthesized phage gene products and induce cell death or dormancy to prevent phage spread^1–4^.

A staggering phage diversity exists in the biosphere, likely driving the requirement for numerous, also extremely diverse immune pathways and concomitant anti-immune mechanisms. The ⌽KZ-like family of jumbo phages (that is, those with genomes >200 kb) possesses many unique attributes including pan-resistance to known DNA-targeting immune systems^5,6^. The ⌽KZ-like phages assemble a proteinaceous nucleus-like structure, where phage genome replication and transcription occur, while phage mRNA is extruded out of the nucleus and translated in the bacterial cytoplasm^7,8^. RNA-targeting CRISPR-Cas systems and engineered nucleases that bypass the nucleus barrier can stop phage propagation, but all DNA-targeting CRISPR systems or restriction endonucleases that have been tested cannot^5,6,9^. RNA-targeting CRISPR systems are relatively rare and are not endogenously present in *Pseudomonas aeruginosa*^10,11^, the host for the best studied jumbophage, ⌽KZ. Therefore, it remains unclear how *P. aeruginosa* and most of their other hosts resist phages of this family.

Here, we identify “jumbophage killer” (Juk), a widespread immune system that specifically detects and blocks ⌽KZ-like jumbophages early in infection. Juk consists of a broadly conserved sensor (JukA, previously YaaW) and a variable effector (JukB), which co-localize at the infected bacterial pole with ejected phage DNA and high copy phage proteins that transit from the phage head into the cell. Successful immunity manifests as a block to early transcription and degradation of the phage genome, thus rescuing the cell by preventing phage-induced bacterial genome degradation and the assembly of phage nucleus.

## Results

### Discovery of an immune system specifically targeting ⌽KZ-like nucleus-forming jumbophages

To identify putative immune systems responsible for resistance to ⌽KZ infection, we infected a panel of 62 *P. aeruginosa* clinical isolates, under liquid infection conditions. We observed that ~50% of the tested isolates were resistant to ⌽KZ infection (Supplementary Fig. 1). Such resistance could be caused either by the absence of bacterial factors, such as receptors, that are required for ⌽KZ infection or by the presence of bacterial immune mechanisms targeting key steps of the ⌽KZ infection cycle. Amongst the resistant strains, model clinical isolate PA14 drew our attention because its grew well across most multiplicities of infection (MOI) but was inhibited by ⌽KZ infection only when MOI > 5 (Figure 1A). This growth inhibition was caused by ⌽KZ phage replication (Figure 1B), which did not occur at lower MOI, whereas the sensitive strain PAO1 was inhibited by ⌽KZ replication at MOIs as low as 10^-6^ (Figure 1A, 1B). These data suggest that PA14 can be permissive for ⌽KZ infection, however, the strong resistance against lower MOIs implies the existence of yet unknown immune mechanisms that can be overwhelmed at high MOI.

**Figure 1:**
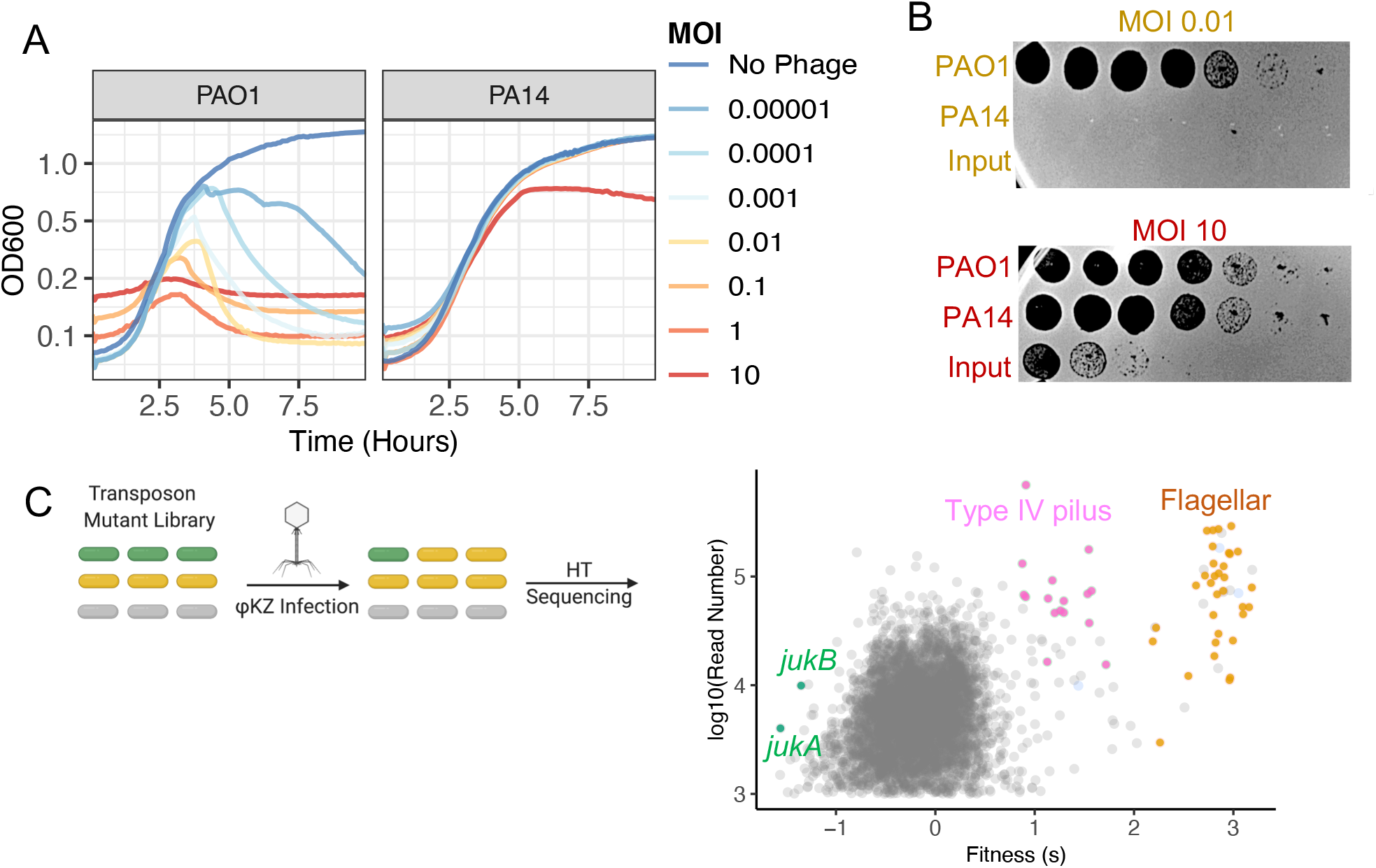
Discovery of the jumbophage killer (Juk) system. **A)** Bacterial growth curves (OD600) of *Pseudomonas aeruginosa* isolate PAO1 or PA14 across a range of multiplicities of infection (MOI). Each line represents ⌽KZ infection at a different MOI. **B)** Titration of ⌽KZ input and output from PAO1 or PA14 infection at MOI 0.01 or MOI 10 (from A) on a lawn of sensitive bacteria (PAO1). **C)** Cartoon schematic of the transposon (Tn) mutant library of PA14 was constructed and used for identification of sensitive or resistant mutants against ⌽KZ infection. Each dot represents a transposon insertion in the PA14 genome and its fitness and read depth are shown after being exposed to ⌽KZ. Genes of interest are highlighted.

To discover immune genes responsible for ⌽KZ resistance in PA14, a pooled PA14 transposon (Tn) mutant library was constructed, infected with ⌽KZ, and subjected to next-generation sequencing to identify transposon insertions (Figure 1C). Mutants with disrupted immune system components were expected to be sensitized to ⌽KZ infection whereas mutants lacking genes required for ⌽KZ propagation would resist infection. Indeed, the Tn screen data (Figure 1C) showed that disruption of flagellar and Type IV pilus genes increased bacterial fitness, suggesting that both structures are required for ⌽KZ infection. Phage adsorption assays confirmed that in *filF:Tn* (flagellum) mutants, phage attachment was abolished (Supplementary Fig. 2A).

Two mutants with Tn insertions in *PA14_03360* and *PA14_03350* showed decreased fitness (that is, increased phage sensitivity) upon ⌽KZ infection (green dots in Figure 1C). *PA14_03360* and *PA14_03350* comprise a predicted two-gene operon; hereafter, we refer to these as jumbophage-killer (Juk) genes *jukA* and *jukB*, respectively. Deletion of either gene sensitized PA14 to ⌽KZ infection (Figure 2A, 2C), which could be complemented *in trans* (Supplementary Fig. 3A), suggesting that both *jukA* and *jukB* are required for immunity against ⌽KZ. When introduced into ⌽KZ-sensitive strain PAO1. *jukA* and *jukB* together, but not either gene alone, conferred resistance against ⌽KZ (Figure 2B, 2C, Supplementary Fig. 3B). When present on a plasmid in PAO1, *jukA-jukB* protected cells from phage-induced lysis more robustly than when in single copy in the chromosome (Figure 2B), likely due to ~10-fold higher mRNA levels (Supplementary Table 1). Collectively, our data demonstrate that the two-gene *juk* operon is necessary and sufficient to provide resistance against ⌽KZ infection.

**Figure 2:**
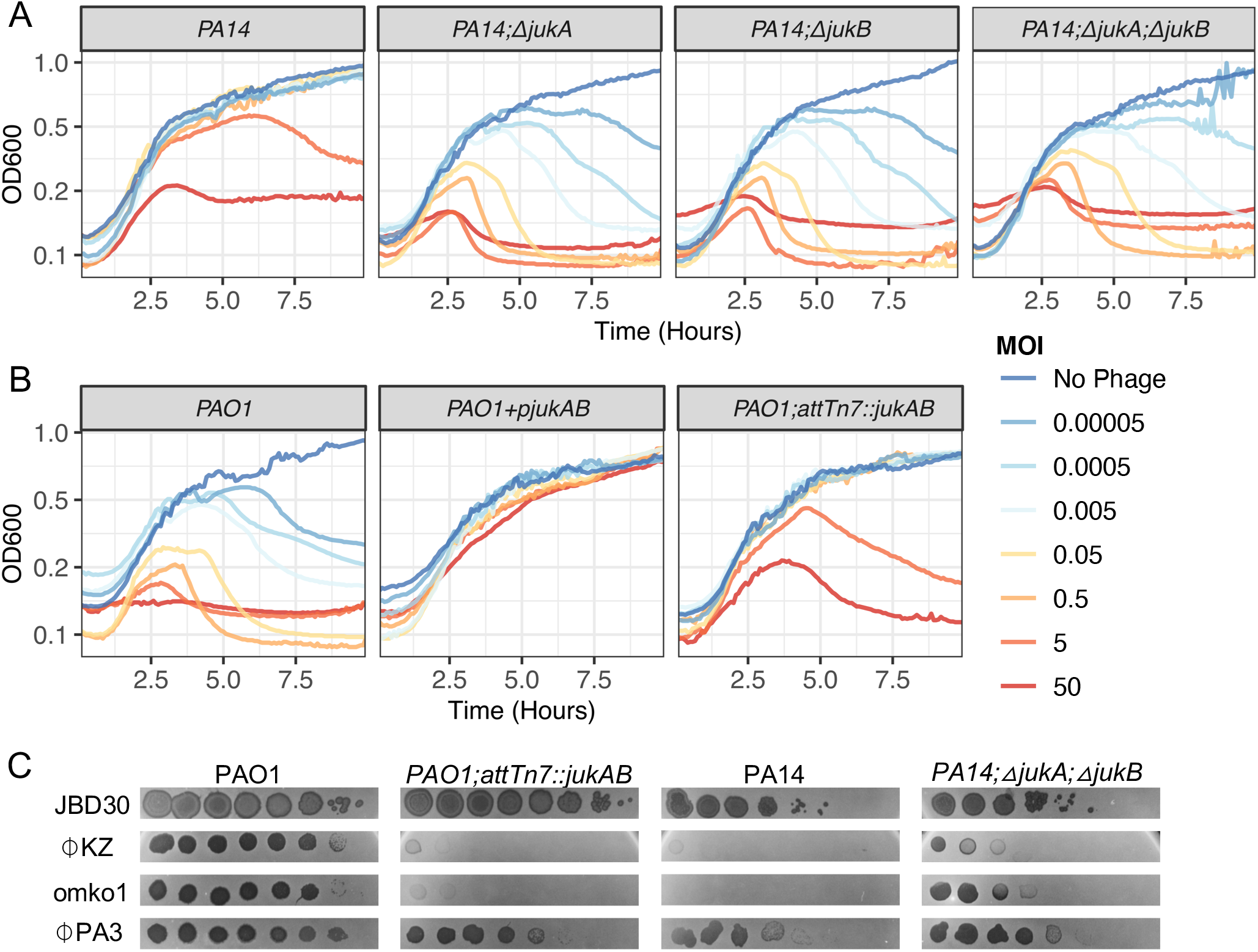
Immune genes *jukA* and *jukB* are necessary and sufficient to provide resistance against nucleus-forming jumbophages. Growth curves measuring OD600 during ⌽KZ infection in **A)** PA14 and indicated mutants or **B)** PAO1 heterologously expressing *jukA* and *jukB (jukAB*) via either plasmid (*PAO1+pjukAB*) or chromosome integration (*PAO1;attTn7::jukAB*). **C)** Phage titration (10-fold serial dilutions) on indicated lawns (more phages tested in Supplementary Figure 4).

To test the specificity of the Juk immune system, we conducted phage infection assays with a panel of phages from a wide range of families (Supplementary Fig. 4, Supplementary Table 2). In addition to restricting the growth of ⌽KZ, Juk also blocked a closely related phage omko1 and displayed weak protection against ⌽PA3 (Figure 2C). Both omko1 and ⌽PA3 belong to the ⌽KZ-related phage family and form a nucleus during infection. By contrast, Juk does not target the unrelated jumbophage PA5oct (Supplementary Fig. 4) or any other phage tested (e.g. JBD30 in Figure 2C and Supplementary Fig. 4). Thus, Juk immunity appears to be specific towards ⌽KZ-related jumbophages.

### Jumbophage killer (Juk) system does not act via known immune mechanisms

PSI-BLAST search identified 1,104 non-redundant JukA homologs in 5,701 genomes (Supplementary Table 3). In eukaryotes, the members of this family are subunits of mitochondrial chaperones involved in the mitochondrial *bc1* cytochrome complex^12^ (e.g. Cbp3, PF03981; HHpred probability 98.6%)., whereas in bacteria, their functions were previously unknown. The JukA family proteins align with Cbp3 throughout their length except for a small N-terminal domain identified in Pfam as DUF3944 (PF13099), but the most pronounced sequence similarity is concentrated within the chaperone domain that facilitates co-translational folding of cytochrome *b*^12^. JukB belongs to a much smaller family with 187 non-redundant proteins in 279 genomes, mostly present in proteobacteria, cyanobacteria and chloroflexi, and has no clearly predicted molecular function (Supplementary Table 3).

With limited functional information available on JukA and JukB, we decided to first test if Juk functions via mechanisms similar to known immune systems. Specifically, we tested whether the Juk system affects phage adsorption, has nuclease activity, or triggers abortive infection. First, PA14 and *PA14ΔjukAB* had similar adsorption kinetics (Supplementary Fig. 2B), although adsorption to this strain, irrespective of Juk immunity, is generally slow. Since this hinders plaquing and microscopy assays, for the rest of the study we use PAO1 and *PAO1;attTn7::jukAB* to study the mechanism of Juk immunity (Supplementary Fig. 2C). Second, JukA and JukB showed no sequence or structural similarity to any known nucleases, and moreover, the multiple alignments of both families did not contain patterns of conserved charged or polar residues known to form unique nuclease active sites, suggesting that Juk is unlikely to directly cleave the ⌽KZ genome. Third, using fluorescence microscopy, infected *PAO1;attTn7::jukAB* cells continued cell division without committing suicide (Supplementary Video 1), indicating that Juk does not cause abortive infection. This is consistent with cell density measurements presented above showing maintained cell viability during phage infection when functional JukA-JukB was over-expressed (*PAO1+pjukAB* in Figure 2B). Thus, Juk most likely targets unique features of the infection cycle of ⌽KZ-related jumbophages via mechanisms not yet identified in other defense systems.

### Juk targets the early stage of ⌽KZ infection

Using fluorescent microscopy, we followed the infection cycle of ⌽KZ to identify which stage of the ⌽KZ infection cycle Juk acts on. ⌽KZ nucleus formation was completely abolished in *PAO1;attTn7:jukAB* strain at MOI 1, whereas a mature ⌽KZ nucleus was observed around 40 minutes post infection in PAO1 (Figure 3A). Furthermore, the ⌽KZ genome created a DAPI-stained puncta upon ejection, which was visible in 60% (n = 306 cells) of PAO1 cells but only 14% (n = 284 cells) of *PAO1;attTn7:jukAB* cells (Figure 3B), suggesting that the genome can enter the cell but then is eliminated. Time-lapse imaging also showed the disappearance of the ⌽KZ genome in the presence of *jukAB* (Supplementary video 2).

**Figure 3:**
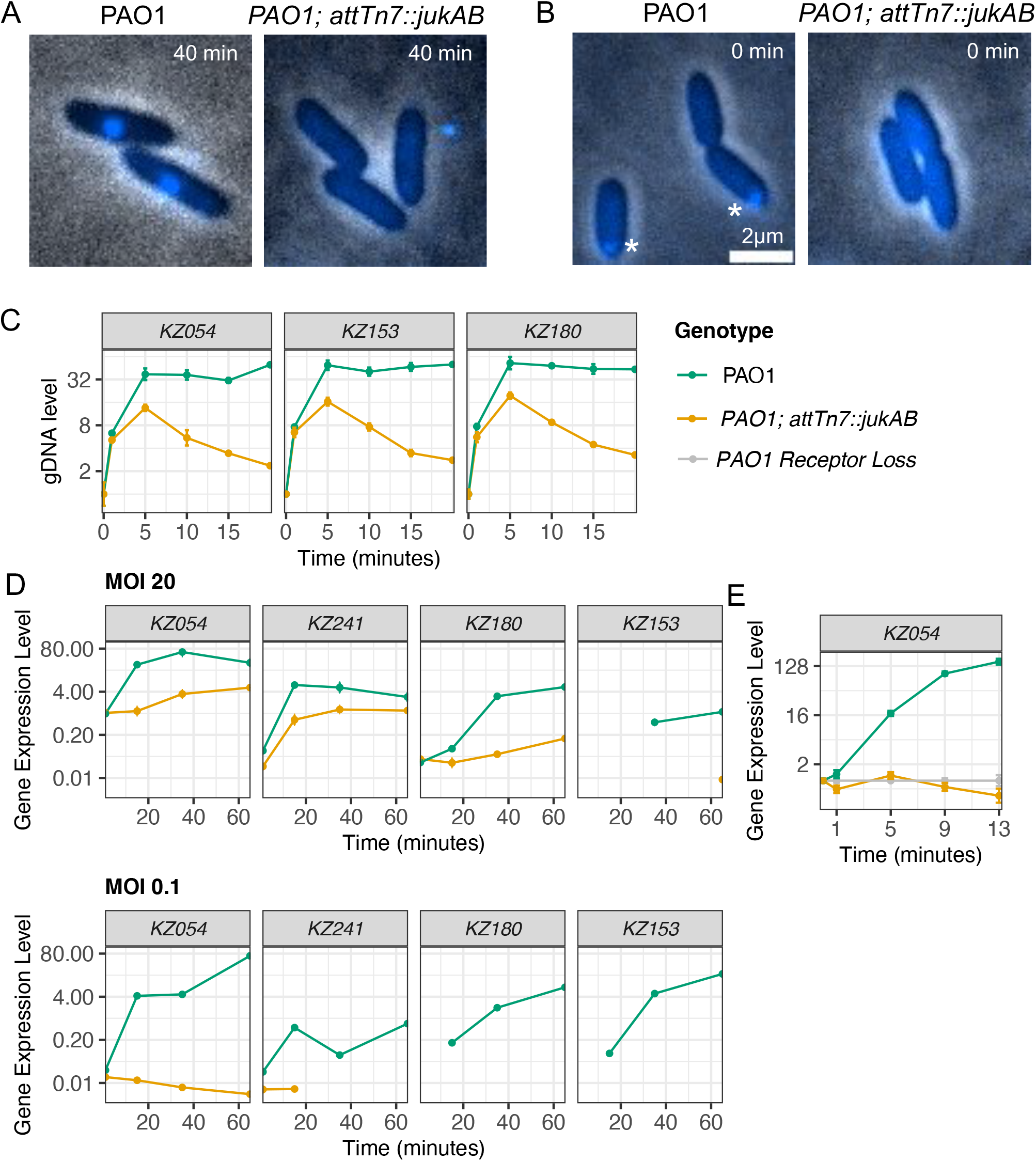
Juk immune system acts on early ⌽KZ infection. ⌽KZ infection of indicated PAO1 cells under DAPI staining at **A)** 40 or **B)** 0 minutes post infection. ⌽KZ genome is marked by *. **C)** Quantification of the DNA level of three ⌽KZ genes over the first 20 minutes of infection (MOI 0.5) in the presence or absence of *jukAB*. **D)** Transcription level of ⌽KZ early (*KZ054* and *KZ241)*, middle (*KZ180*) and late (*KZ153*) genes at MOI 20 or 0.1. Points below the assay detection limit were eliminated. **E)** Quantification of the expression level of *KZ054* during early infections at MOI 0.2. Note that ⌽KZ is unable to inject its genome into the receptor loss mutant. Error bars in **C)**, **D)**, and **E)** represent one standard deviation calculated from two technical replicates.

To corroborate the microscopy observations, we measured phage gDNA levels at three distinct loci (*KZ054, KZ153*, and *KZ180*) by quantitative PCR (qPCR). Upon infection at MOI 0.5, ⌽KZ gDNA level initially increased in both PAO1 and *PAO1;attTn7:jukAB*, from 0 to 5 minutes, suggesting that ⌽KZ adsorption and DNA ejection occurred at this stage (Figure 3C). After 5 minutes, ⌽KZ DNA level remained stable in PAO1 prior to rapid amplification, but decreased in *PAO1;attTn7:jukAB* (Figure 3C, Supplementary Fig. 5). The rate of ⌽KZ gDNA decrease in *PAO1;attTn7:jukAB* (~8 fold from 5 min to 20 min, Figure 3C) was much faster than the rate of cell division (~30 min per cell division), suggesting that the amount of ⌽KZ gDNA dropped not solely due to dilution caused by cell division, but rather, as a result of gDNA degradation.

During infection, ejected ⌽KZ RNA polymerase immediately starts the transcription of early ⌽KZ genes^13^. To confirm the action of Juk on the early stages of ⌽KZ infection, we quantified the expression levels of two early genes, *KZ054* and *KZ241*, one middle gene, *KZ180*, and one late gene, *KZ153*^13^, at MOI 0.1, where Juk successfully neutralizes ⌽KZ infection and at MOI 20, where ⌽KZ overwhelms Juk immunity and completes its infection cycle. At MOI 0.1, all four genes were effectively silenced by Juk, whereas at MOI 20, Juk immunity only partially lowered transcript levels (Figure 3D). More extensive sampling of early gene *KZ054*, which encodes the major protein of the phage nucleus^7,8^, revealed that expression was completely inhibited in *PAO1;attTn7:jukAB* at MOI 0.2, whereas it increased ~100-fold in PAO1 from time 0 to 5 minutes (Figure 3E). Together, these findings suggest that Juk immune proteins act immediately post infection to halt the phage transcription, subsequently leading to indirect ⌽KZ DNA degradation, presumably by host nucleases.

### JukA is the infection sensor in the two-component Juk immune system

To examine how Juk immunity defends against ⌽KZ infection, we fluorescently tagged JukA and JukB (*PAO1; attTn7::sfCherry2-jukA; jukB-mNeonGreen*) and followed their localization. Fluorescent fusion did not affect Juk immunity (Supplementary Fig. 3C). Without ⌽KZ infection, JukA was diffuse in the cytoplasm whereas JukB appeared as motile puncta (Figure 4A, Supplementary Fig. 6A). However, upon ⌽KZ infection, JukA and JukB rapidly clustered at cell poles where the phage infection occurred (Figure 4B, Supplementary Fig. 6B). As discussed above, the ⌽KZ genome is often rapidly cleared, but in cells where a DAPI-stained puncta (that is, ejected phage DNA) could be visualized (Arrow in Figure 4B), JukAB co-localized with it (Figure 4B). To identify the driver of this polar localization phenotype, each protein was expressed on its own, showing that JukA sensed the infection on its own and localized to the infection site (Figure 4C), whereas JukB puncta formation and localization were entirely JukA-dependent (Figure 4D). Because JukA by itself is not sufficient to abrogate infection, the co-localization of JukA puncta and the ejected ⌽KZ DNA was more apparent in the absence of JukB (Figure 4C). These findings suggest that JukA serves as a ⌽KZ sensor, which recruits JukB.

**Figure 4:**
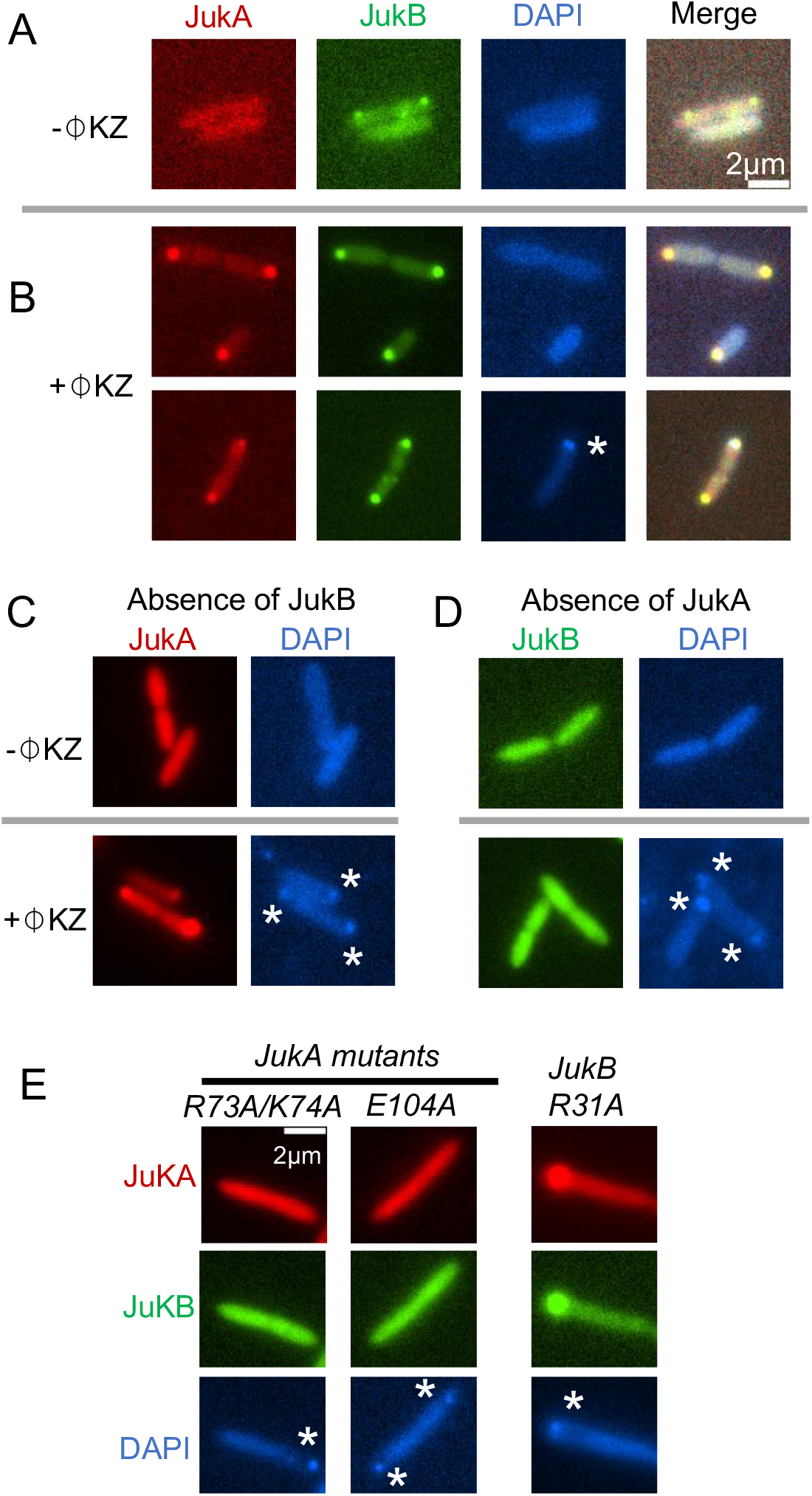
JukA and JukB localize to the infected cell pole. In each image, sfCherry2-JukA, mNeonGreen-JukB, and DAPI stained DNA are shown. Ejected ⌽KZ genome is marked by *. JukA and JukB localization **A)** without and **B)** with ⌽KZ infection. **C)** JukA localization in the absence of JukB. **D)** JukB localization in the absence of JukA. **E)** Localization of JukA and JukB proteins in *jukA* and *jukB* loss of function mutants.

To further investigate the role of the Juk localization in resisting ⌽KZ infection, we mutated conserved amino acids in JukA and JukB based on multiple sequence alignment of Juk homologs from several *Pseudomonas* species (Supplementary Fig. 7A, B). Five of the 14 JukA mutants resulted in complete loss of immune function (Supplementary Fig. 7C) and all five abolished JukA localization (Figure 4E). In contrast, one JukB mutation (R31A) led to the complete loss of Juk immunity (Supplementary Figure 7D) but did not impact its localization during ⌽KZ infection (Figure 4E). Thus, this site in JukB is likely involved in a downstream interaction or catalytic activity.

### Juk activity is triggered by ejected ⌽KZ factors

While ⌽KZ-like phages ⌽PA3 and omko1 induced JukA polar localization (Supplementary Fig. 8A), small dsDNA phages DMS3 and JBD30, which also infect at the pole via the Type IV pilus, did not (Supplementary Fig. 8A). Thus, JukA is likely triggered by factors specific to ⌽KZ-like jumbophages. To determine whether ejected ⌽KZ factors or newly synthesized proteins induce the JukA response, cells were treated with translation inhibitor gentamicin 5 minutes prior to ⌽KZ infection. Phage infection in untreated cells proceeded normally (Figure 5A), whereas gentamicin blocked infection progression and nucleus formation (Figure 5B). We observed that cells treated with gentamicin maintained the typical JukA and JukB localization to the infection pole (Figure 5C and D), suggesting that Juk activity is triggered by ejected ⌽KZ factors.

**Figure 5:**
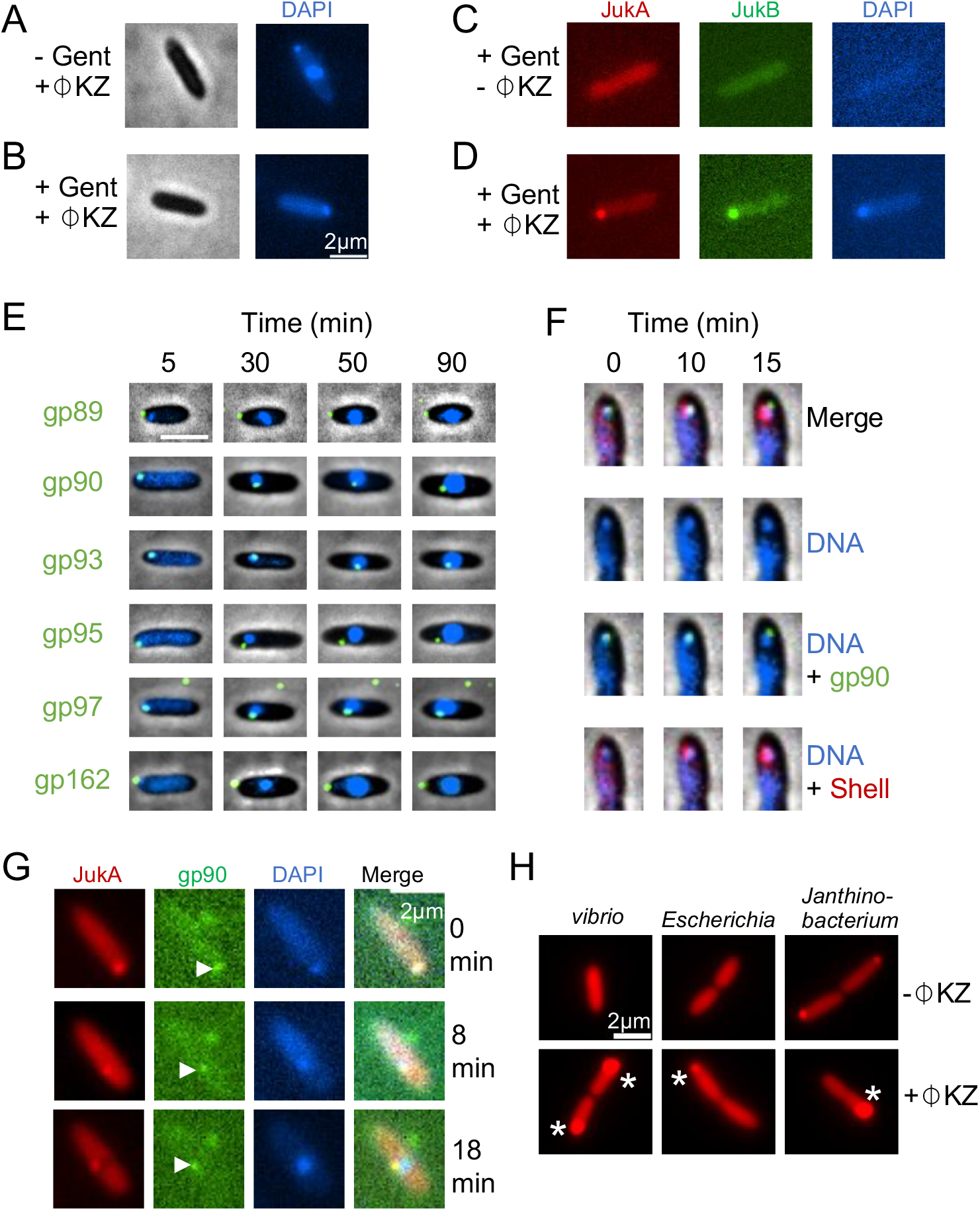
Juk activity is triggered by ejected ⌽KZ proteins. PAO1 cells infected in the **A)** absence or **B)** presence of gentamicin (Gent). PAO1 cells expressing sfCherry2-JukA and mNeonGreen-JukB **C)** uninfected or **D)** ⌽KZ infected in the presence of gentamicin. **E)** The localization of ejected IB proteins (mNeonGreen tagged) over the course of infection. **F)** Localization of ⌽KZ DNA (blue, DAPI-stained), ejected IB protein (mNeonGreen tagged), and the assembling nascent nucleus (mCherry tagged “shell” protein). **G)** sfCherry2 tagged JukA imaged along with ejected gp90 (tagged by mNeonGreen; markd by arrowhead). DAPI staining was applied in **A)-G)** to reveal the phage genome. **H)** Localization of sfCherry2 tagged JukA homologs from the single-component Juk immune systems. JukA puncta are marked by *.

The ⌽KZ phage head packages both RNA polymerase^13^ and a large proteinaceous “inner body” (IB) structure around which the genome is wrapped^14^. Abundant virion proteins that likely make up this structure have been identified via mass spectrometry^15^, but their specific function(s) remain unknown. We hypothesized that abundant IB proteins could be ejected into host cells, potentially triggering Juk immunity. We fluorescently tagged each IB protein (⌽KZ gp89, gp90, gp93, gp95, gp97, gp162) with mNeonGreen (Supplementary Fig. 8B) by infecting PAO1 expressing each IB-mNeonGreen fusion with wild-type ⌽KZ. As a result, the newly assembled phages packaged a mixture of wild-type and labeled IB proteins in their heads, forming individual fluorescent foci visible under the microscope (Supplementary Fig. 8B). During infection, gp90, gp93, gp95, and gp97 were ejected along with ⌽KZ DNA, whereas gp89 and gp162 appeared not to enter the host, forming a green focus that remained on the cell surface (Figure 5E). Fluorescent labeling of the phage nucleus protein gp54 (mCherry-gp54) clearly showed the phage DNA being “handed off”, without initiating replication, from the ejected protein-DNA complex to the nascent nucleus at ~10-15 minutes after infection (Figure 5F). From that time point on, gp90, gp93, and gp97 remained tightly associated with phage nucleus throughout the intracellular phage development, whereas gp95 frequently dissociated (Figure 5E). These observations demonstrate the ejection of four ⌽KZ inner body proteins into the host cell during infection, while two remain in the virion.

We then tested whether JukA co-localized with the ejected IB proteins by infecting a bacterial strain expressing sfCherry2-JukA alone with ⌽KZ virions containing fluorescent tagged IB proteins. The ⌽KZ genome, gp90, and JukA localized together at the beginning of infection. As the nucleus assembly progressed, however, the ⌽KZ genome dissociated from JukA and gp90, whereas JukA and gp90 remained co-localized (Figure 5G). Thus, most likely, JukA recognizes ejected IB protein(s) or other ejected proteins rather than the ejected ⌽KZ genome itself. However, overexpression of single IB proteins in the host was insufficient to trigger the re-localization of Juk proteins (Supplementary Fig. 8C). Furthermore, we constructed *△KZ093* and *△KZ097* knockout phages (these genes encode gp93 and gp97, respectively), but these mutants still triggered Juk immunity as indicated by microscopy (Supplementary Fig. 8A) and no spontaneous phage escape mutants were obtained despite many efforts. Interestingly, *KZ090* and *KZ095* appear to be essential for phage reproduction as knockouts could not be obtained. Thus, JukA likely recognizes an essential IB protein or a higher order complex.

### JukA sensor is a component of numerous predicted immune systems in diverse bacteria

Gene context analysis of *jukA* genes revealed a strong link with defense systems (Supplementary Table 3, Supplementary Fig. 9A). At least one previously characterized or predicted defense gene is present in 173 of 329 representative loci (see Supplementary Fig. 9A for a phylogenetic tree made from these 329 JukA homologs). WYL domain^16,17^ and components of restriction-modification systems are most frequent defense associated genes identified in *jukA* neighborhoods. Apart from *jukB* (present in 46 of representative loci), several other gene families are expected to form an operon with jukA. In particular, these include O-antigen ligase RfaL, which affects adsorption rates in *Salmonella*^18^. Yet uncharacterized PD-(D/E)XK and HNH family nuclease domain containing proteins, Xre family transcriptional regulators are also found in vicinity of *jukA* genes. Several uncharacterized proteins, such as DUF2541 (PF10807) and EcsC-like (PF12787), form putative operons and are likely coexpressed with *jukA* (Supplementary Figure 9B). Furthermore, JukA fusions to Ras-like GTPase, patatin-like phospholipase and a domain of unknown function containing a coiled coil region were also identified (Supplementary Figure 9C). Based on all these observations, we speculated that JukA might function as a jumbo phage sensor that combines with various effectors.

17 operons covering different parts of the phylogenetic tree were synthesized, transformed into PAO1, and tested against a panel of phages (Supplementary Table 4). Eight of these 17 operons contained *jukB* as the putative effector, originating from *Leclecia, Jinshanibacter, Desulfolutivibrio, Flammevirga, Stenotrophomonas, Rhodospirillum, Shewanella*, and *Vibrio*. Note that the JukB homologs from *Shewanella* and *Vibrio* have low sequence identity with the originally identified JukB. Seven of the eight JukAB homologs blocked ⌽KZ replication, like the original Juk system from PA14 (Supplementary Fig. 10 and 11). JukAB from *Flammevirga* was the only one that did not provide immunity against any phage assayed. Deletion of either *jukA* or *jukB* from the functional *Jinshanibacter, Stenotrophomonas*, *Rhodospirillum or Shewanella jukAB* operons abolished the immune function, indicating that these distinct Juk variants are also two-component systems (Supplementary Fig. 11). Surprisingly, JukA from *Vibrio* alone is sufficient to provide immunity against ⌽KZ (Supplementary Fig. 11).

Nine predicted operons with diverse putative effectors (Supplementary Fig. 9A; Supplementary Table 4) were next assayed. Two operons from *Escherichia* and *Janthinobacterium*, provided specific immunity against ⌽KZ-related jumbophages (Supplementary Fig. 10 and 11). *Escherichia* and *Janthinobacterium jukA*s are paired with Xre family transcription regulators. Deletion of either *jukA* or the putative partner gene showed that in both cases, JukA was again sufficient for immunity (Supplementary Fig. 11). Even though *Escherichia, Janthinobacterium* and *Vibrio* JukA proteins appear sufficient for immunity, they contain no identifiable additional domains. To test whether these putative single protein immune systems block ⌽KZ infection similarly to JukAB, we tagged them with sfCherry2 and observed that these JukAs have the same subcellular distribution as the original JukA sensor from PA14, namely, diffuse in cell without phage infection but rapidly concentrating to the infection site upon phage infection (Figure 5H). However, unlike original Juk immunity, where the phage genome disappears over time, these JukA proteins appear to arrest ⌽KZ infection at early stage without subsequent phage genome degradation (Supplementary Video 3), suggesting a different immune mechanism. The existence of functional anti-⌽KZ immunity in diverse bacteria suggests that these bacteria are likely subject to infection by ⌽KZ-like jumbophages. ⌽KZ-like jumbophages so far have been discovered in few families of bacteria. These results suggest that this phage family is more widespread than currently observed.

## Discussion

Well-characterized intracellular defense systems in bacteria recognize either the ejected nucleic acids, replication intermediates, or synthesized phage proteins^3^. For example, CRISPR-Cas and restriction-modification systems typically target phage DNA rapidly in the cell prior to replication, whereas other, recently described anti-phage systems, such as DarTG^19^ and Nhi^20^, appear to detect replicating DNA. In contrast, the recently described Avs (AntiVirus STAND)^21^, CapRel systems^22^, and likely CBASS^23^ and Pycsar^24^, detect late expressed phage structural proteins. However, to our knowledge, no mechanism that detects ejected phage proteins has been reported. Here, we propose that the “jumbo phage killer” (Juk) system detects ejected phage factors to enable specific and robust non-abortive immunity (Figure 6). The sensor protein JukA localizes to the phage infected pole as the first stage of immune activation, which recruits effector JukB and prevents phage progression (Figure 6). JukA localization does not rely on de novo synthesis of phage proteins, and intriguingly, overlaps with the localization of ejected inner body (IB) proteins, supporting the mechanistic connection between ejected factors and immunity (Figure 6). In addition to the highly abundant IB proteins, ⌽KZ-like phages also package a virion RNA polymerase complex and likely other low copy number proteins in the head^15^. It remains formally possible that any other ejected factor(s) trigger JukA and the specific identity of its interaction partner(s) remains to be determined. We consider phage DNA itself to be an unlikely trigger due to the lack of any obvious DNA binding or nuclease domains in JukA or JukB, the apparent sequestration of this phage genome throughout infection, and the continued co-localization of JukA with IB proteins after the genome moves to the assembled nucleus.

**Figure 6:**
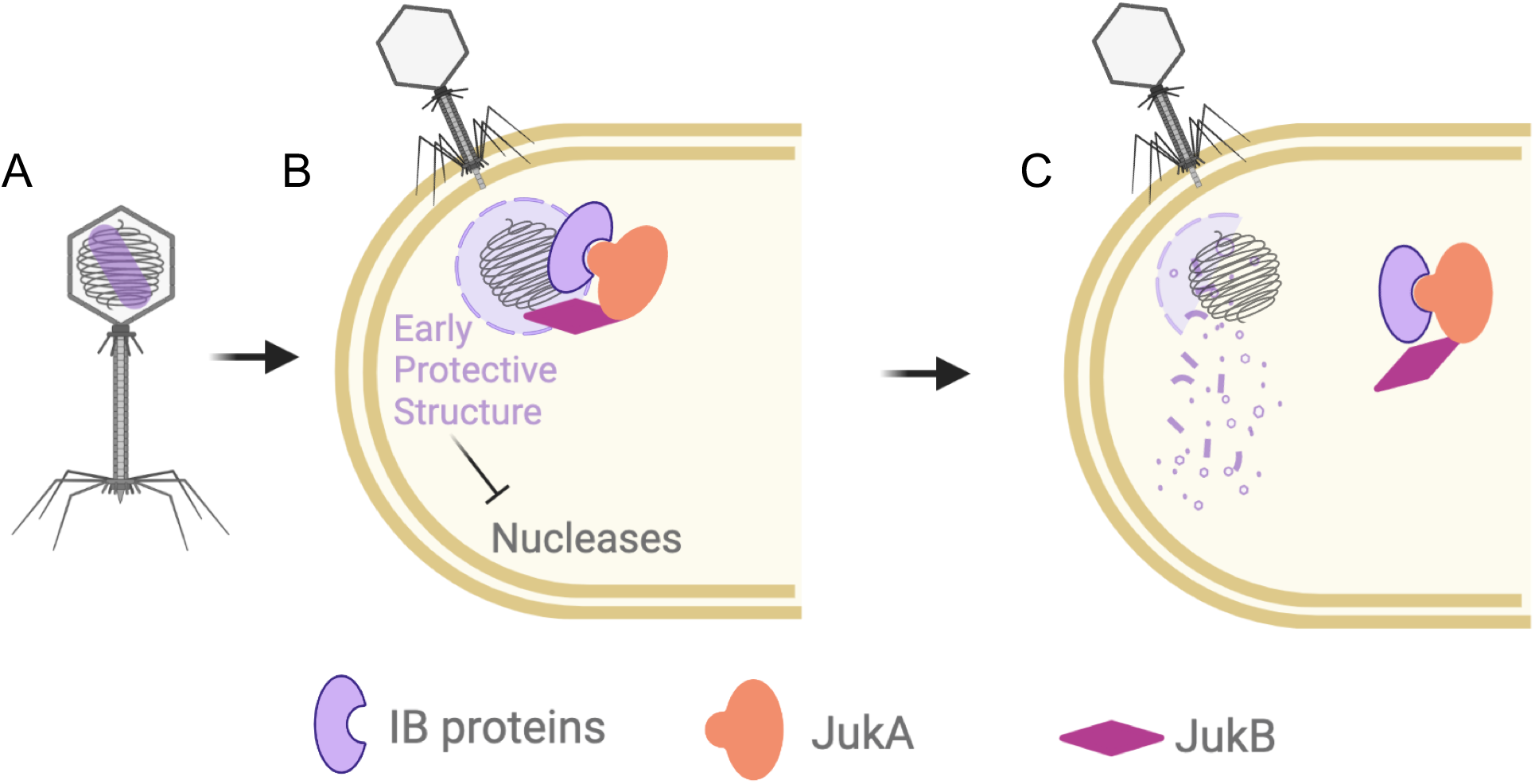
Proposed mechanistic model for Juk immunity. **A)** ⌽KZ virion has phage genome spooled around the ”rod-like” inner body (IB) protein structure. **B)** Upon infection, sensor JukA recognizes ejected IB proteins and are recruited to the phage infection site. JukA further recruits the effector JukB. **C)** JukB facilitates the destabilization of the early protective structure. The interactions among phage IB proteins, JukA, and JukB remain in cells.

The immunity provided by Juk and its homologs appears to be specific to ⌽KZ-like jumbophages, which are so far unique in their ability to assemble a proteinaceous nucleus that protects phage genome from restriction and CRISPR nucleases. Given that purified phage DNA is sensitive to such nucleases in vitro, an early structure must protect the phage genome before the nucleus forms. Indeed, we previously reported that experimentally arrested infections left a stable DNA puncta at the pole, with no degradation of phage DNA by restriction enzymes^5^. This protective structure could be the “round compartments”^25^ or “unidentified spherical bodies”^26^ revealed by recent electron microscopy studies. The experiments reported here identified some of the likely IB protein components of this structure and uncovered a novel immune strategy that apparently destabilizes it, rendering the phage DNA susceptible to cytoplasmic nucleases. Without Juk immunity, some of the ejected IB proteins play an essential role, initiating an intimate “hand-off” of the phage genome to the assembling nucleus.

JukA is a widespread bacterial protein whereas the spread of JukB is far narrower. We observed that JukA formed putative operons with potential effectors distinct from JukB and furthermore found that some JukA proteins provided the anti-phage protective effect on their own although recruitment of additional components *in trans* cannot be ruled out. Adding credence to this possibility, we identified cases where JukA and JukB were encoded in different genomic locations (Supplement Table 3). These findings demonstrate functional versatility and modularity of the Juk system, which is a hallmark of microbial defense systems^3,4,27^. Furthermore, the widespread nature of Juk seems to suggest that nucleus-forming jumbophages targeted by JukA are more common than presently appreciated.

There are many questions regarding the specific mechanisms of Juk anti-phage protection. Nevertheless, it seems clear that Juk represents a distinct modality of anti-phage defense whereby phage proteins ejected into the host cell are targeted before any phage specific process, such as expression of immediate early genes, takes place. Such mechanisms might be advantageous compared to those targeting late phage proteins in that they not only abrogate the phage reproduction but also allow the infected cell to survive, without the need to induce dormancy or programmed cell death as is the case with the Avs systems^21^ and some CRISPR variants^28^. This work generates the potential that targeting ejected phage components, for example, the early transcription apparatus, is a common defense strategy, in which case many such systems, beyond Juk, remain to be identified.

## Supporting information

Supplementary Table 1, 2, 4 and more

## Acknowledgements

This work in the Bondy-Denomy lab was supported by the National Institutes of Health (NIH) (R01GM127489, R01AI171041-01) and the UCSF Program for Breakthrough Biomedical Research funded in part by the Sandler Foundation. K.S.M. and E.V.K. are funded through the Intramural Research Program of the National Institutes of Health of the USA (National Library of Medicine). DAA is funded by NIH (R35GM118099). We thank Dr. Stephen Lory for sharing the strains and plasmids used to construct Tn mutant library and Dr. Michael Boucher for sharing expertise on Tn genomic library construction. We thank Dr. Andrea Fossati and Bondy-Denomy lab members for constructive comments and suggestions

## Author contributions

Y.L. conceived the project, designed and performed all experiments, and wrote the manuscript. J.G. contributed to microscopy and inner body labeling. S.H. contributed to execution of experiments. E.C. contributed to generate the whole genome sequencing data. D.A.A. contributed to experiments design. K.S.M. and E.V.K executed the bioinformatic analysis. J.B.D. supervised the project, designed experiments, and wrote the manuscript.

## Methods and Materials

### Bacterial growth and transformation

The strains, plasmids, and phages used in this study are listed in the Supplementary Table. P. aeruginosa strain PAO1 and PA14 was grown in LB at 37 °C with aeration at 300 rpm. When necessary, plating was performed on LB agar with carbenicillin (250 μg/ml) or gentamicin (50 μg/ml). Gene expression was induced by the addition of l-arabinose (0.1% final).

Plasmids are delivered into PAO1 with electroporation, while conjugation between PA14 and plasmid-carrying E. coli SM10λpir is used to deliver plasmids into PA14. The conjugated cell mixture containing PA14 and SM10 was streaked out on Vogel-Bonner minimal medium (VBMM) agar containing antibiotics. VBMM media selects against *E. coli* cells and antibiotics selects against PA14 cells that do not contain the desired plasmid^29^.

### Genetic manipulation

For chromosomal insertions at the attTn7 locus, PAO1 cells were electroporated with the integrating vector pUC18T-lac and the transposase expressing helper plasmid pTNS3, and selected on gentamicin. Potential integrants were screened by colony PCR with primers PTn7R and PglmS-down. Electrocompetent cell preparations, transformations, integrations, selections, plasmid curing and FLP-recombinase-mediated marker excision with pFLP were performed as described previously^30^.

Two-step allelic exchange was used to delete *jukA, jukB*, or *jukA&B* from the PA14 genome, Empty vector pMQ30 was used to construct allelic exchange vectors via Gibson Assembly. The allelic exchange vectors were then transformed into PA14 via conjugation. After first-crossover, which occurs shortly after conjugation, VBMM agar containing 50 μg/mL gentamicin was used to select for PA14 merodiploids. Subsequently, sucrose counter-selection was used to select for double crossovers. Sucrose-resistant colonies were then subject to gentamicin to select for sucrose-resistant and antibiotic-sensitive colonies as successful outcomes. The desired colonies are further confirmed via PCR amplification and Sanger sequencing. The protocol is detailed in^29^.

### Phage plaque assays and phage propagation

100 μl of appropriate overnight culture was suspended in 3 ml of 0.45% molten top agar and then poured onto an LB agar plate containing 10 mM MgSO4 and appropriate antibiotics. After 10-15 min at room temperature, 2 μl of ten-fold serial dilutions of phages was spotted onto the solidified top agar. Plates were incubated overnight at 37°C.

For high-titre phage lysates, PAO1 overnight culture was diluted 100-fold in Fresh LB media and infected with phages at MOI 0.01. Phages were collected after overnight infection. Phage stocks were stored at 4 °C and used for routine infection assays.

### Growth curve experiments

Growth curve experiments were carried out in a Synergy H1 microplate reader (BioTek, with Gen5 software). Cells were diluted 1:100 from a saturated overnight culture with 10 mM MgSO4 and antibiotics and inducers, as appropriate. Diluted culture (140 μl) was added together with 10 μl of phage to wells in a 96-well plate. This plate was cultured with maximum double orbital rotation at 37 °C for 24h with OD600 nm measurements every 5 minutes.

### PA14 Transposon mutant library Screening

#### Construction of PA14 transposon (Tn) mutant library

Transposon insertion mutants were generated by mating PA14 and E. coli SM10λpir carrying the suicide vector pBTK30^31^. The mini-transposon pBTK30 is a suicide delivery vector (ori R6K) that contains a mariner C9 transposase, an origin of transfer (oriT RK2), and a β-lactamase gene (bla) specifying ampicillin resistance. The 1.5-kilobase transposable element is located between 28 base pair inverted repeats and consists of an aacC1 gene (providing gentamicin resistance) that is transcribed toward a transcriptional and translational terminator. Successfully conjugated PA14 cells were selected for on VBMM agar containing 50 μg/ml gentamicin. The selected PA14 cells contain Tn insertions at random sites in the genome. ~200,000 mutant colonies were collected and pooled together. The pool of PA14 Tn mutant library was aliquoted and stored at −80 °C at 10^^^10 cells/ml.

#### Phage treatment

~10^9 PA14 Tn mutant cells (~10,000 cells/mutant) were removed from −80 °C, diluted 2-fold into fresh LB media containing 50 μg/ml gentamicin and 10 mM MgSO4, and recovered at 37 °C, 300 rpm for 2 hours. The recovered mutant library was then split into 6X 100 μl replicates with two replicates treated with ⌽KZ at MOI 30 as infection samples, two treated with SM buffer as non-infection controls, and the rest two collected as samples at time 0. The infection samples and non-infection controls were immediately grown at 37 °C and 300 rpm in a 96-well plate with their OD600 measurements being monitored. Cells were collected via centrifugation after ~5 hours of treatment.

#### Genome extraction and library preparation

Genome extraction was conducted using Qiagen Deasy UltraClean Microbial Kit. ~1 μg genomic DNA from each sample was used to construct the sequencing library using NEB Illumina Library Prep Kit. Customized primers carrying unique multiplexing tags and multiplexing indices were designed and used for adaptor ligation and library amplification. Primer YL001 was annealed to YL002 or YL003 and used as the adaptor in the adaptor ligation step. The downstream transposon junctions were amplified using a two-step PCR protocol with the second step as a nested PCR reaction to reduce nonspecific PCR amplification.

#### Sequencing and analysis

DNA libraries were sequenced using Illumina next-seq technology with >2 million reads per sample. Sequencing data were trimmed using the software Cutadapt^32^ and aligned to the reference genome (NC002516.2) using Bowtie^33^. Transposon junctions were extracted. The number of reads for each transposon junctions was counted. Assuming that transposon insertions within the same gene shared similar phenotypes, we treated mutations within the same gene as a mutant group, referred to the mutant group as “mutant” hereafter. Note that only mutants with Tn insertions in the coding regions were considered in our analysis.

Next, to evaluate the effect of bacterial genes in ⌽KZ infection, we compared the frequency of mutants in the presence and absence of ⌽KZ infection and calculated their fitness using the equation: s = ln(‘MutantFreq_W/_⌽KZ’ / ‘MutantFreq_W/O_⌽KZ’). If a gene is important in ⌽KZ resistance, disrupting this gene should make bacterial more susceptible to ⌽KZ infection, leading to a lower mutant frequency in the presence of ⌽KZ infection and thus resulting in a negative fitness estimate.

The majority of mutants did not affect ⌽KZ resistance/sensitivity and had a fitness centered around 0. The fitness distribution of neutral mutants is roughly Gaussian where mutants with a low read number heavily contribute to the left and right tail of the Gaussian distribution. Mutants outside of the Gaussian distribution are likely to be “non-neutral” mutants that affect ⌽KZ infection.

### Adsorption assay

Adsorption assay were conducted by infecting exponentially growing bacteria with phage at MOI 0.01 in a flask. Infected bacteria cultures were grown at 37 °C with gentle shaking at 60-80 rpm. For each timepoint, 50 μl of samples were removed from the flask and added to 450 μl of SM buffer containing extra chloroform. Samples were then centrifuged at 5000X g for 5 minutes and the supernatants were used for plaque assays and virion quantification.

### Fluorescence microscopy and imaging

#### Agarose pad preparation

LB containing 10 mM MgSO4 will be referred to as LBM. 0.064 gram of agarose were added into a mixture of 2 ml LBM and 6 ml H2O, melted and kept at 55 °C. DAPI was added to the melted gel liquid to reach a final concentration of 0.5 μg/ml. Pour the gel liquid onto assembled slides to form agar pads.

#### Cell preparation

Dilute overnight bacterial culture 100-fold into fresh LBM and grow cells at 37 °C with aeration at 300 rpm until reaching OD600 ~0.4. If phage infection is needed, mix cell culture and phages to a desired MOI and incubate in a dry block at 30 °C for 10 minutes. Pipette 1 μl of bacterial cell onto a piece of agarose pad and assembled onto slides for imaging. Note that *jukB* expression is ~80 fold higher than the chromosomal integrated operon when being expressed on the pHERD30T plasmid and induced by 0.1% arabinose, which leads to JukB aggregation. To avoid artificial protein aggregation, arabinose was not added when imaging cells that contained plasmids expressing *jukA* and *jukB*. The leaky expression was confirmed to be functional.

#### Imaging

Microscopy was performed on an inverted epifluorescence (Ti2-E, Nikon, Tokyo, Japan) equipped with the Perfect Focus System (PFS) and a Photometrics Prime 95B 25-mm camera. Image acquisition and processing were performed using Nikon Elements AR software.

### qPCR and RT-qPCR

Bacteria was grown to the log phase with an OD600 measurement of ~0.4 and infected with ⌽KZ to desired MOIs. 500 μl of cell culture were removed at each timepoint. Cell pellets were immediately collected by spinning down the cell culture at 6000X g. For RNA extraction, cell pellets were flash-frozen using liquid nitrogen and stored at −80 °C. For DNA extraction, cell pellets were stored at −20 °C directly. Total RNA was extracted from the resulting cell pellets by performing acidic phenol-chloroform extractions. Luna® Universal One-Step RT-qPCR Kit from NEB was used for RT-qPCR reactions. Genomic DNA was extracted using standard phenol-chloroform DNA extraction protocol. PerfeCTa® SYBR® Green SuperMix from QuintaraBio was used for qPCR reactions. Both qPCR and RT-qPCR reactions were performed on CFX Connect thermocycler from Bio-Rad.

*Pseudomonas aeruginosa* housing-keeping gene *rpoD* was used as the internal control during calculation. PAO1 receptor loss mutant was not included in all batches of experiments. For the ones that included the receptor loss mutant, gene fold changes were normalized against the readout for the mutant.

### Gentamicin treatment

Gentamicin was added to exponentially growing bacteria (OD600 ~0.4) to a final concentration of 50 μg/ml. Cells were treated with gentamicin at 5 minutes prior to phage infection or microscopy unless it is noted otherwise.

### JukA and JukB mutagenesis

Around the world PCR was used to introduce mutations to *jukA* or *jukB* with PCR primers carrying the desired mutations. Plasmid *pAB04+sfCherry2-jukA;jukB-mNeonGreen* was used as PCR template.

### ⌽KZ virions packaged with fluorescence tagged IB proteins

Each inner body proteins was tagged with mNeonGreen on pHERD30T plasmids. WT ΦKZ was used to infect PAO1 containing each IB-mNeonGreen fusion. As a result, the newly assembled phages packaged a mixture of both wild-type and labeled IB proteins in their heads, forming individual fluorescent foci under the microscope. In contrast, no fluorescent phage was obtained by expressing mNeonGreen alone.

### ⌽KZ genetic editing

Deletion of IB genes from ⌽KZ were performed by following the protocol in^34^.

### Choice of *jukA*-containing operons for immune function assay

*jukA* homologs and their ten neighboring genes (5 upstream and 5 downstream) on bacterial genomes were identified. First, we removed genes that were not in the same operon as *jukA* homologs. Genes in the same operon satisfied two relax criteria: a) genes are on the same strand as *jukA*, and b) genes are within the 50bp away from *jukA* homologs. Second, we used the combination of gene names in the same operon as a unique ID to separate these *jukA* containing operons into different groups. For groups with >8 bacterial genomes, a representative was manually selected. Based on the cost to synthesize the operon and the identity of the putative effector, we eventually chose 17 operons for synthesis and cloning.

### Multiple sequence alignment of Juk proteins

Multiple sequence alignments for JukA and JukB proteins within *Pseudomonas* were performed using Clustal-Omega with its default setting.

### Bioinformatic analysis of the distribution of JukA and JukB homologs

#### Identification of JukA and JukB homologs

PSI-BLAST^35^ search (e-value cut-off was set to 1e-04, three iterations, the rest of the parameters remained default) was performed using JukA protein (WP_003137196.1) as a query against a database of complete Refseq genomes (November 2021 release)^36^. The set was further manually refined: a few false positives were discarded, and few false negatives (JukB neighbors) were included in the final set. Same procedure was applied for identification of JukB homologs.

#### Phylogenetic analyses

JukA sequences were clustered using MMseqs2^37^ with the similarity threshold of 90% identity, and one representative was taken from each cluster for further analysis. Sequences were aligned using a previously described iterative procedure^11^. Based on this alignment, N- and C-terminal domains fused to some of JukA homologs were removed and several short sequences were discarded. The remaining sequences were realigned using the same method. The resulting multiple alignment was further filtered to retain the positions with less than 50% of gaps and homogeneity value greater than 0.1. Approximate maximum likelihood phylogenetic trees for the filtered alignments were built using FastTree (WAG evolutionary model, gamma distributed site rates)^38^. The same program was used to obtain support values.

#### Genomic neighborhood analysis

For each representative *jukA* and *jukB* gene used for phylogenetic analysis, 5 genes upstream and downstream were collected and all proteins encoded in these neighborhoods were annotated using PSI-BLAST^35^ with E-value threshold = 0.01 run against position-specific scoring matrices (PSSMs) deposited in the CDD database^39^. Only hits to regularly updated databases, namely Pfam, CDD and COGs were considered. HHpred search with default parameters against PDB, Pfam and CDD profile databases was used for unannotated proteins or domains^40^. Additionally, all proteins in the respective neighborhoods were clustered using MMseqs2 program^37^ with the similarity threshold of 0.5, and a cluster identifier was assigned for each ORF in the neighborhood. Defense function was assigned based on CDD annotation and a collection of known and predicted defense system components^3,27,41^ or on the presence of *jukB* genes.

---

**Supplementary Figure 1:**
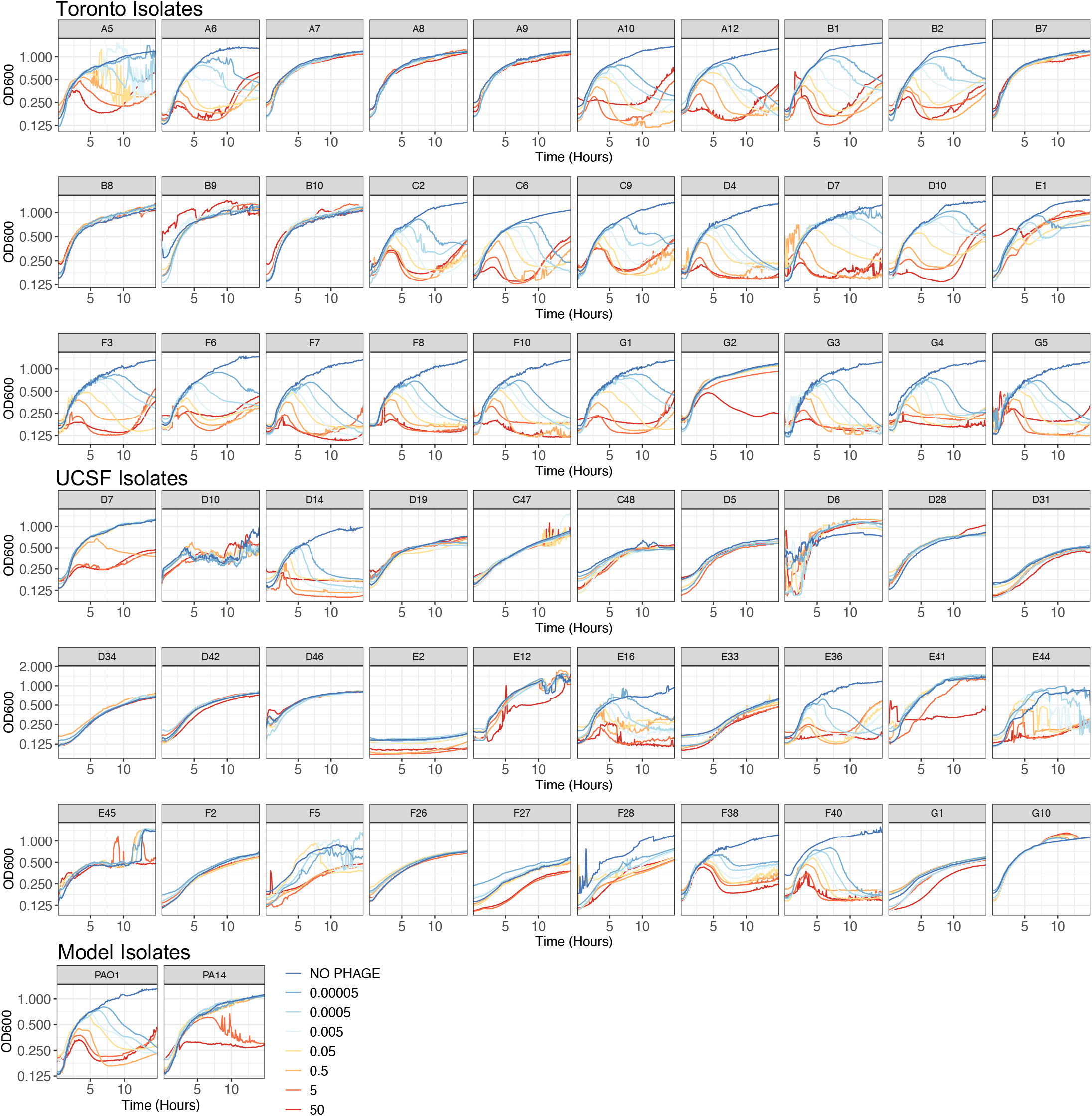
Growth curves (measuring OD600) of a panel of *Pseudomonas aeruginosa* isolates when being infected with phage ⌽KZ at different multiplicities of infection.

**Supplementary Figure 2:**
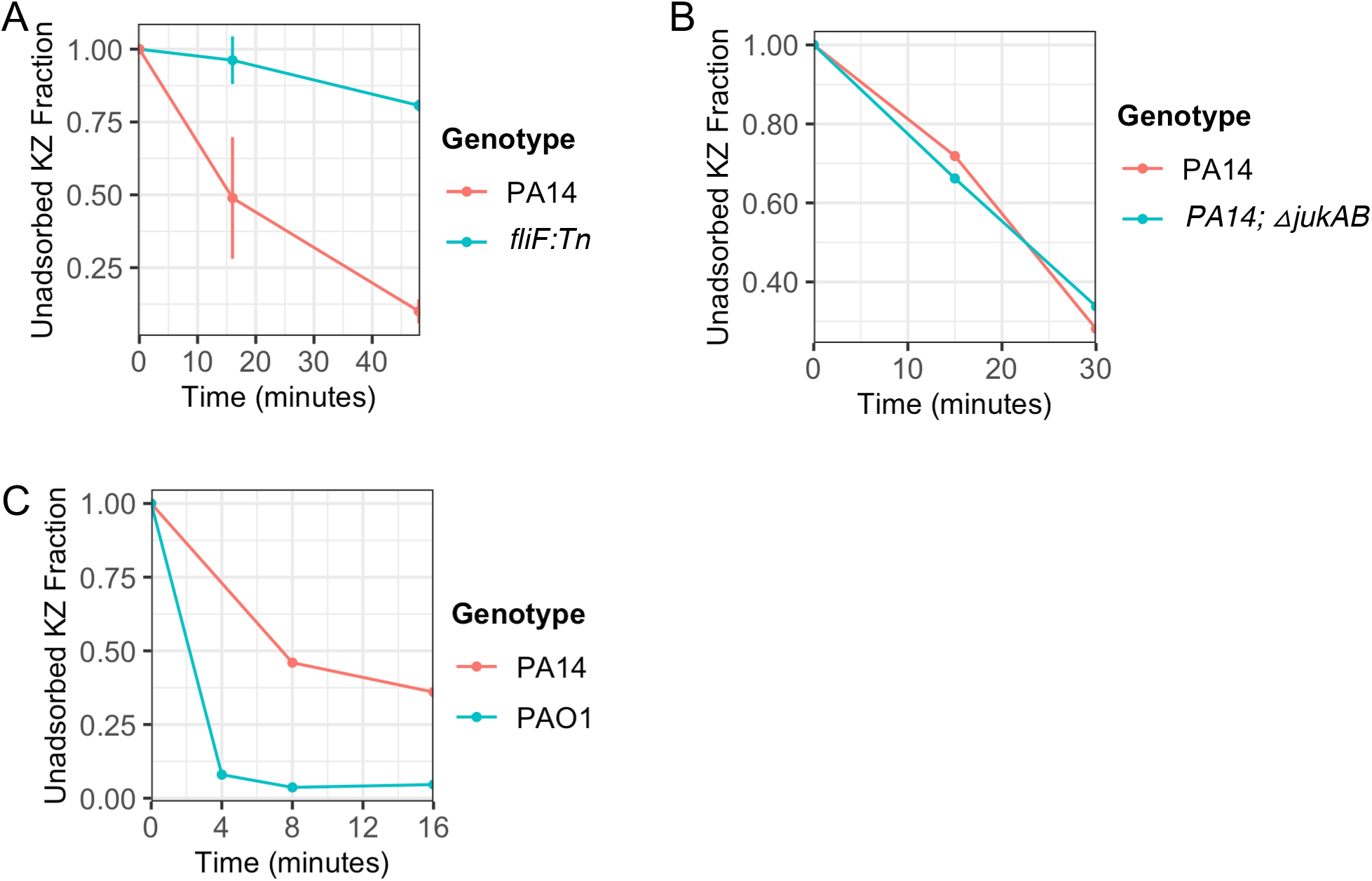
⌽KZ adsorption efficiency in various *P. aeruginosa* strains. The fraction of un-adsorbed phage in the supernatant was calculated by plating on a sensitive strain and plotted as a function of time. Error bar in **A)** represents one standard deviation inferred from two biological replicates. Plot **B)** and **C)** contain one replicate.

**Supplementary Figure 3:**
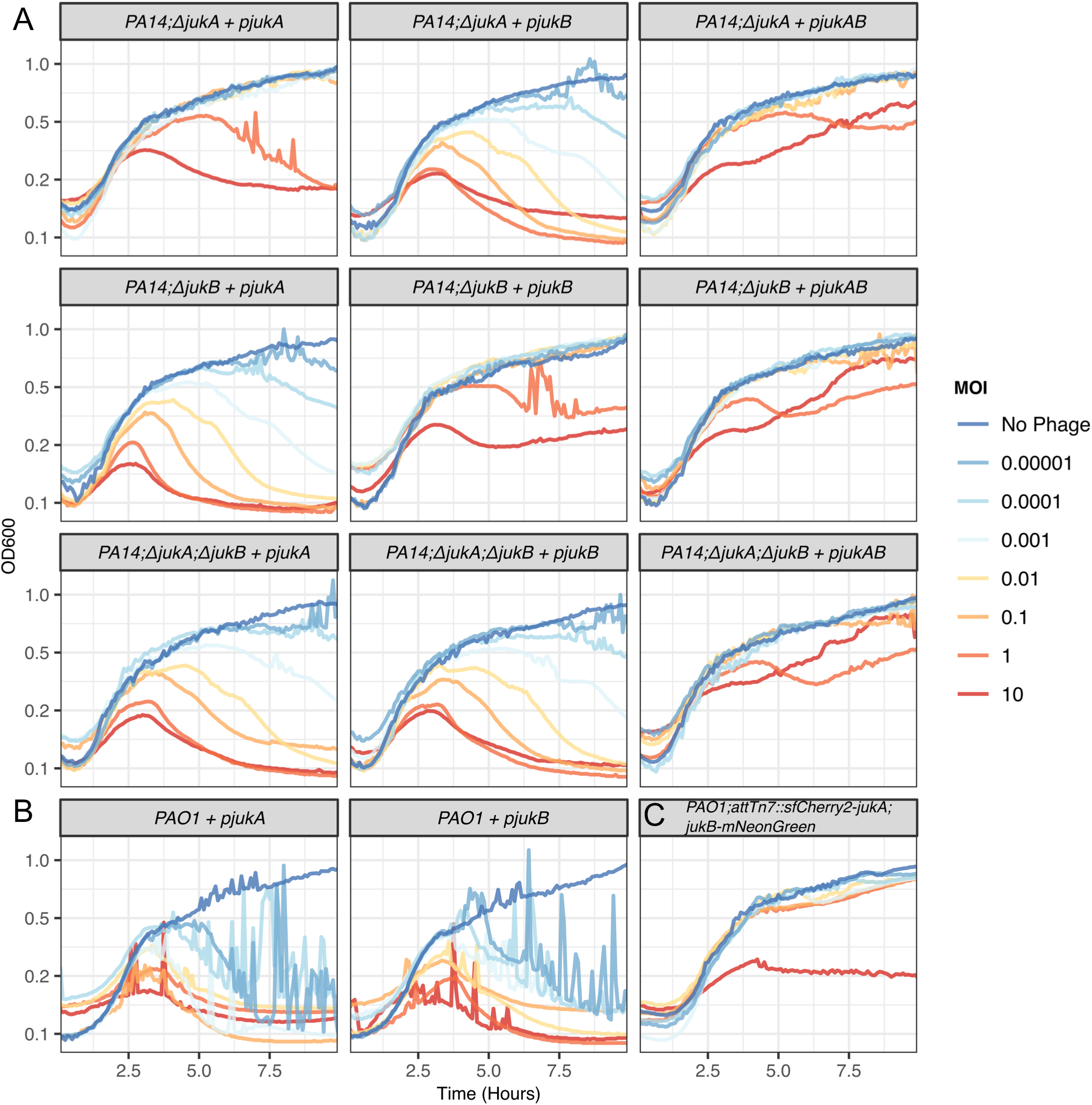
Growth curves (OD600 measured) of indicated strains during infection with ⌽KZ across a range of MOIs. **A)** PA14 deletion mutants are complemented by expression of indicated genes in *trans*. **B)** The indicated constructs are also expressed in PAO1. **C)** Growth curves of PAO1 carrying fluorescence tagged Juk proteins during infection with ⌽KZ across a range of MOIs.

**Supplementary Figure 4:**
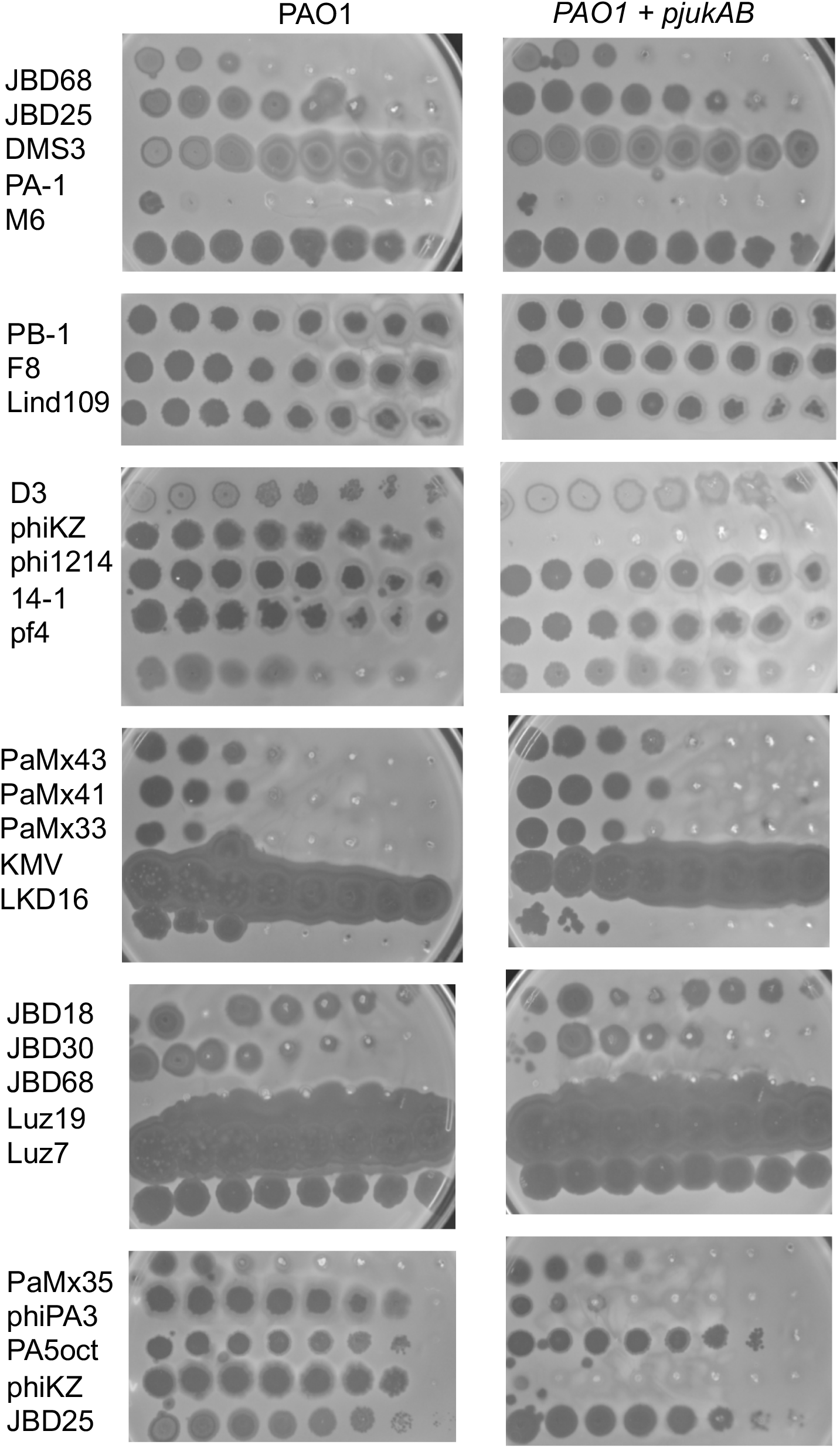
Phage titration assay with 10-fold phage dilutions from left to right, plated on lawns of PAO1 and PAO1 *+ pjukAB*.

**Supplementary Figure 5:**
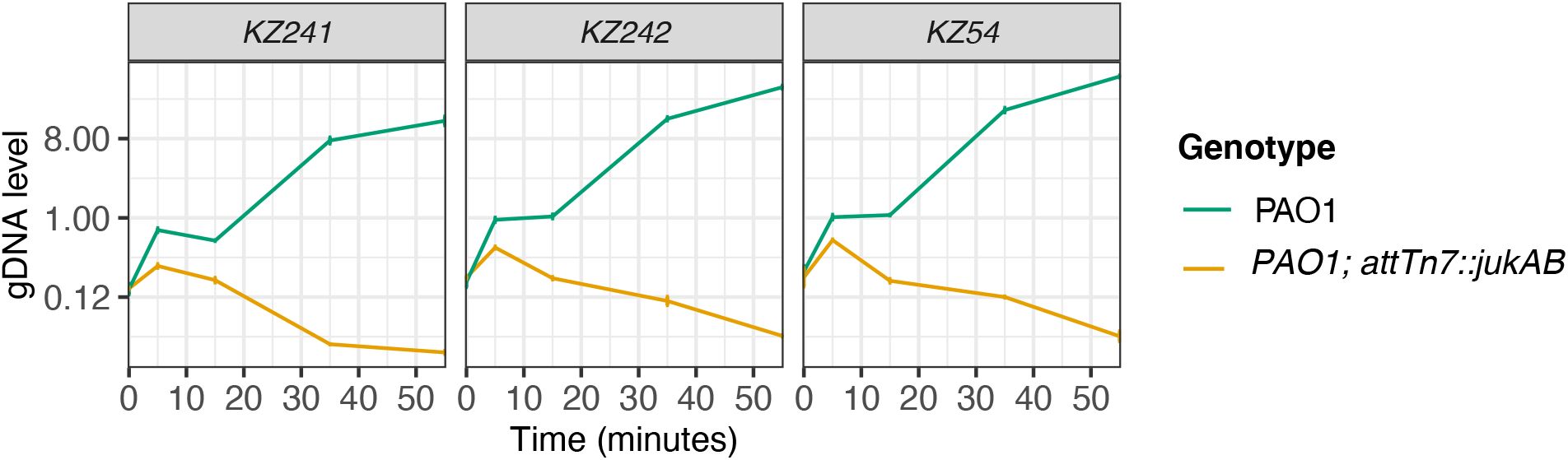
The genomic DNA level of three ⌽KZ genes over the course of infection. Error bar represents one standard deviation inferred from two technical replicates.

**Supplementary Figure 6:**
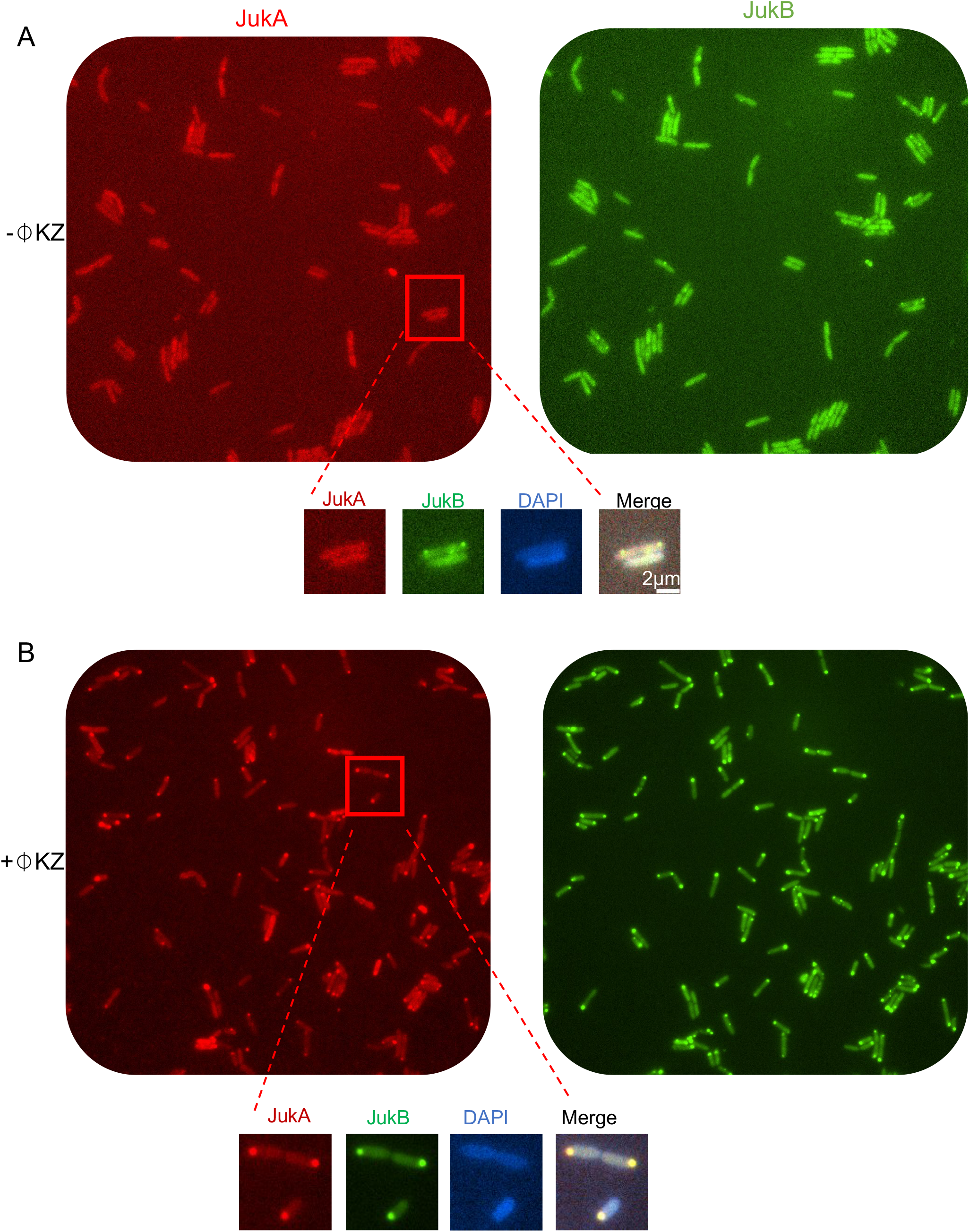
Fluorescence microscopy data of JukA and JukB co-localization at the ⌽KZ infection site. **A)** No phage added. **B)** With ⌽KZ infection.

**Supplementary Figure 7:**
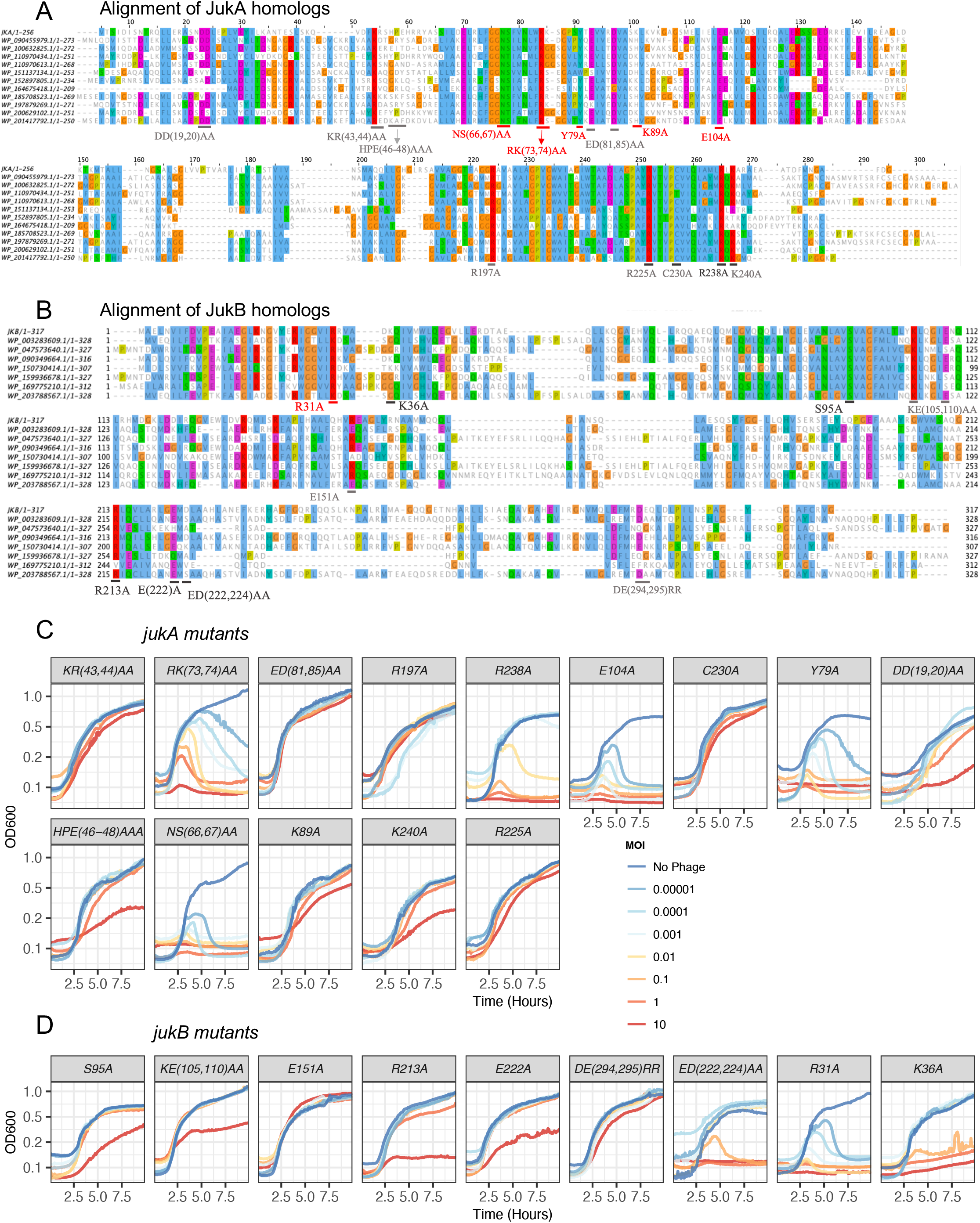
**A,B)** Multiple sequence alignment of **A)** JukA and **B)** JukB homologs. Amino acid residues that were mutated are labeled. Mutations that cause loss of function are in red color. **C, D)** Growth curves OD600 during phage infection of PAO1 expressing *jukAB* with the indicated *jukA* or *jukB* mutants.

**Supplementary Figure 8:**
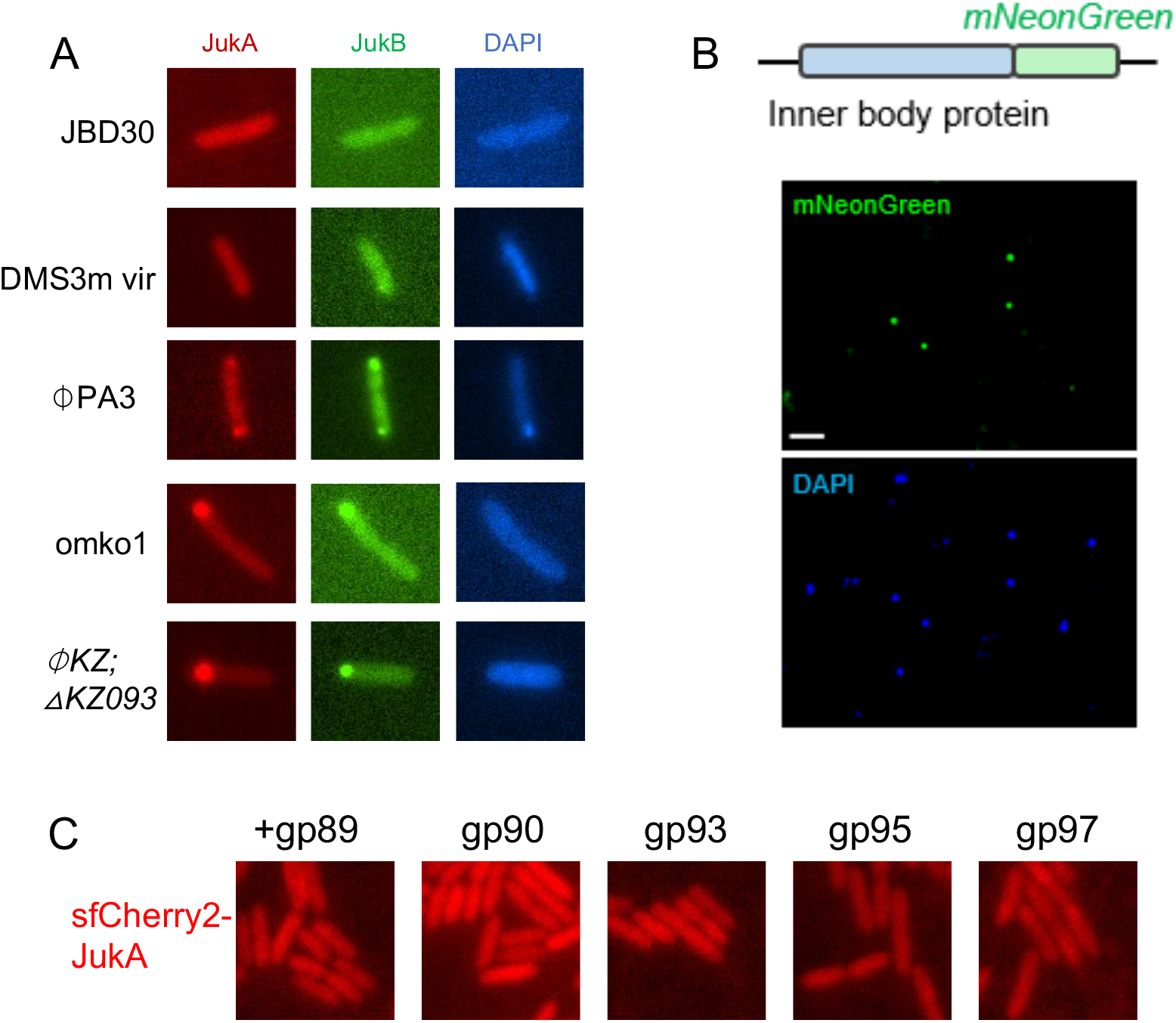
**A)** The localization of Juk proteins when cells are infected by different phages. **B)** Fluorescence microscopy image of ⌽KZ virions containing mNeonGreen tagged inner-body proteins. **C)** Distribution of JukA in cells overexpressing IB proteins with 0.25% arabinose induction.

**Supplementary Figure 9:.**
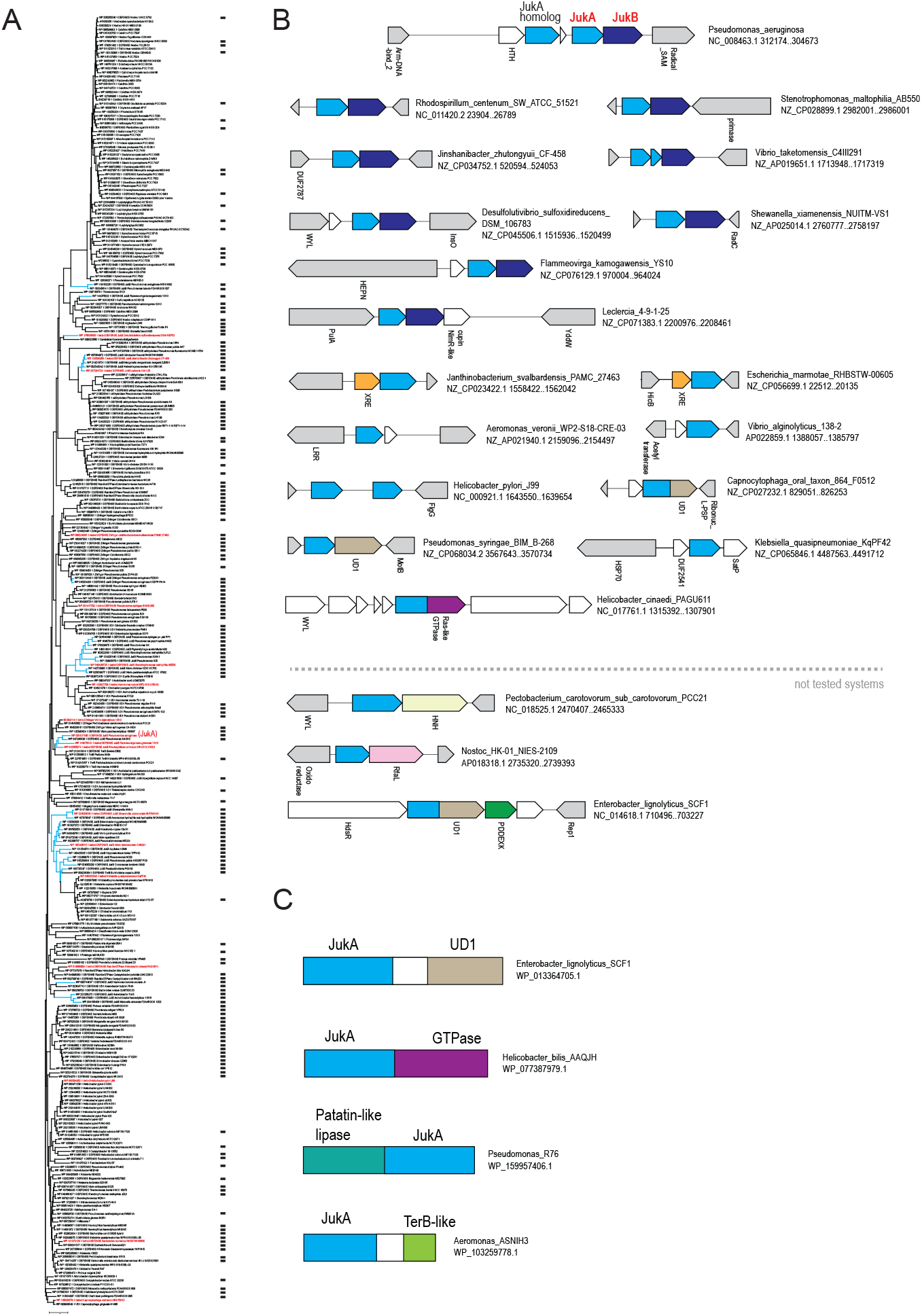
Phylogenetic, gene context and sequence analysis of JukA family. **A)** Phylogenetic tree for 329 representatives of JukA family was built using Fasttree program as described in Methods. The branches leading to JukB present in the same locus as respective JukA are colored blue. Each leaf is denoted by protein identifier, species name and additional information as follows: DEFENSE – at least one known or predicted defense gene is encoded in the respective locus; JukB – JukB is encoded in respective locus; a/b hydrolase (patatin homolog), Ras-like GTPase, TerB, Zn-finger – if respective domains are fused to JukA; UD1 – is an unknown domain either fussed or encoded in the respective JukA locus. Presence of defense genes is also shown by black rectangles on the right. JukA proteins that were experimental tested are highlighted by red. **B)** Organization of *jukA* neighborhoods. Genes are shown as arrows. Genes and untranslated regions are proportional for their size. Species name, nucleotide accession and coordinates of respective region are indicated on the right. Homologous genes or domains are shown by the same color. Only genes or domains that are often associated with j*ukA* are colored. White arrows – genes that do not have any annotation. Gray arrows - flanking genes. Short names for genes or protein families are indicated below the arrows if annotation is available. Abbreviations for short names: WYL – is ligand binding domain of WYL family, often fused to a DNA binding domain; HEPN – protein containing predicted ribonuclease of HEPN family; UD1 – unknown domain 1; HTH – helix-turn-helix, XRE - helix-turn-helix of xre family, HNH and PD-DExK are DNA nuclease of respective families. **C)** Domains fused to JukA. Domains are shown as rectangles roughly proportional to domain size and color coded the same way as in the panel B. Respective protein accession and species name are indicated on the right.

**Supplementary Figure 10:**
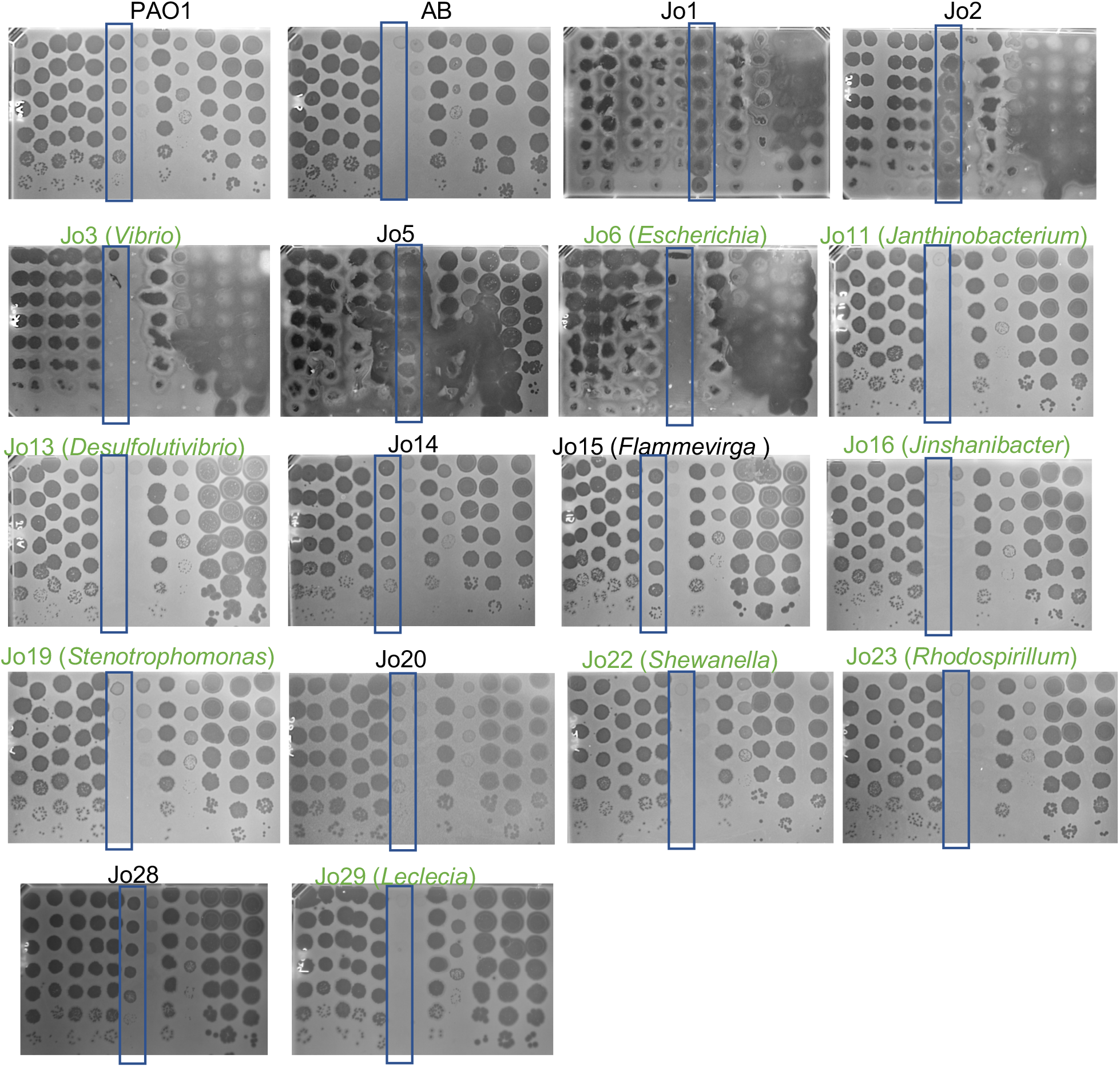
Immunity function of different *jukA*-containing operons when expressed in PAO1. Spot titration plaque assays with 10-fold dilutions from top to bottom on lawns expressing “Juk Operons (Jo)” from various bacteria. Operons that have immune function are labeled in green color. From left to right, the infecting phages are PB-1, 14-1, F8, ⌽1214, Lind109, ⌽KZ, EL, ⌽PA3, PA5oct, M6, YuA and PA-1. phiKZ is highlighted in the rectangle.

**Supplementary Figure 11:**
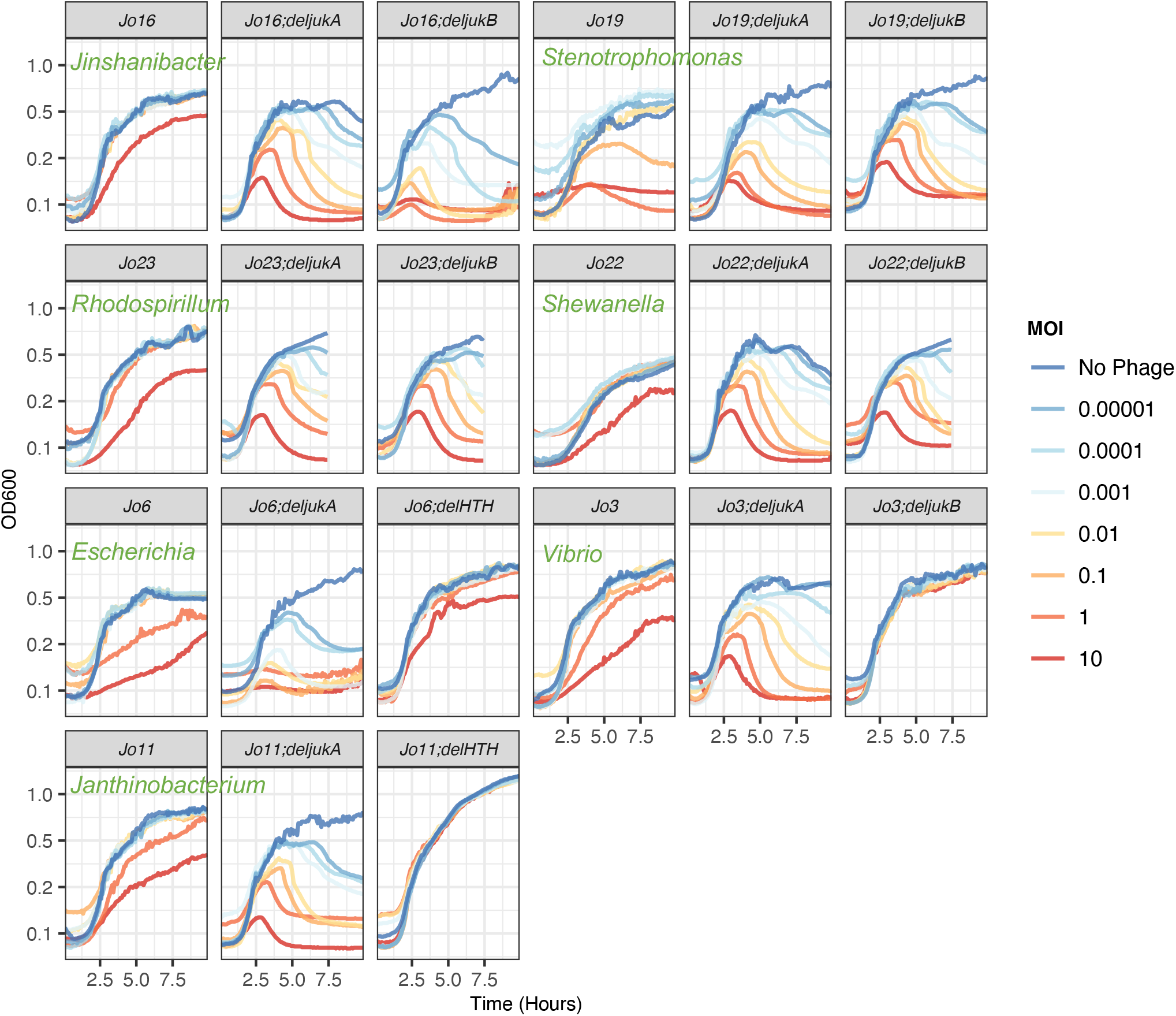
Bacterial growth curves measuring OD600 during phage⌽KZ infection at different MOIs. The indicated ”Juk operons (Jo)” are expressed in PAO1 with the indicated gene deleted. Necessity test of genes in *jukA*-containing operons for immune function against ⌽KZ.

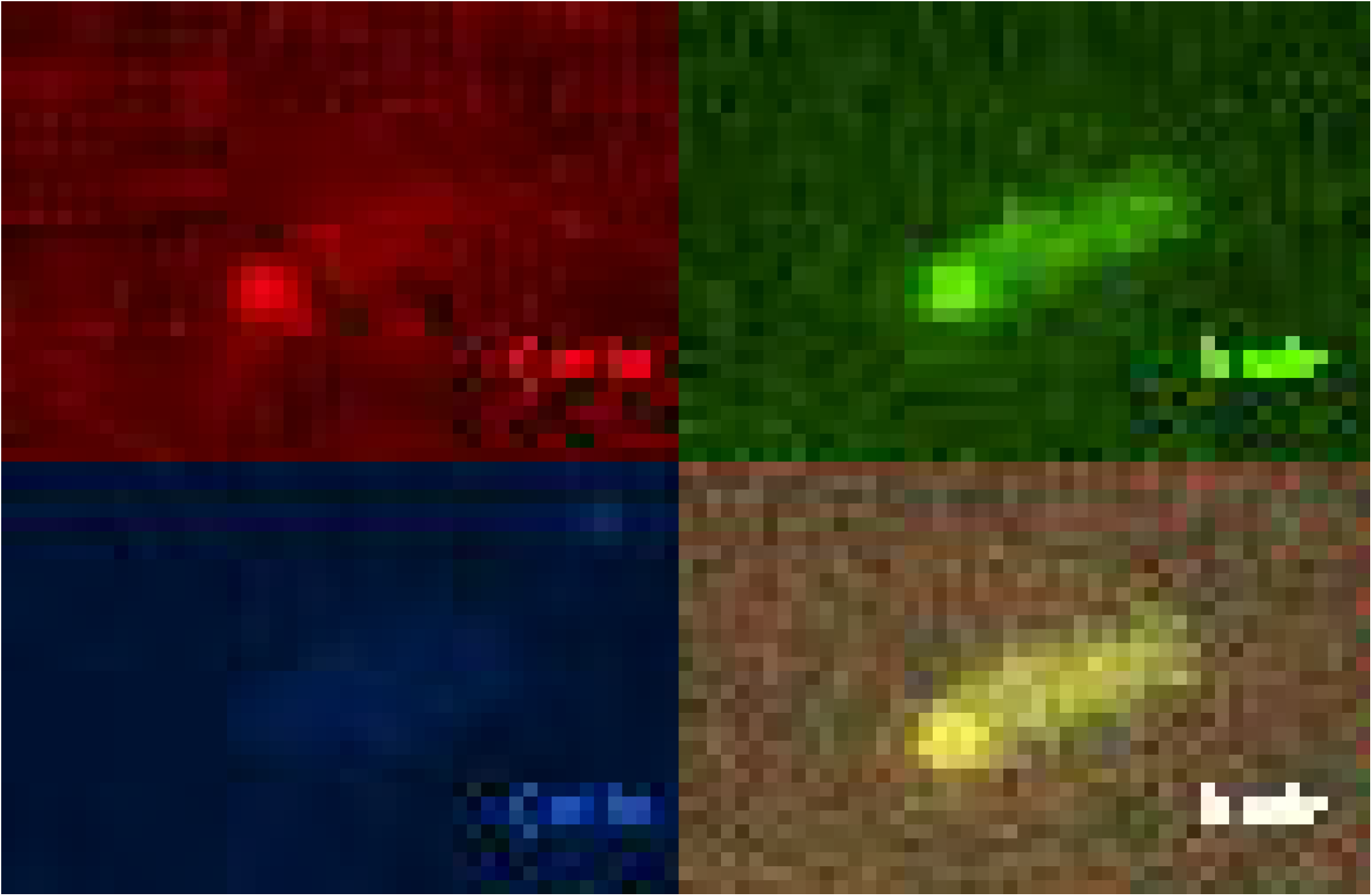

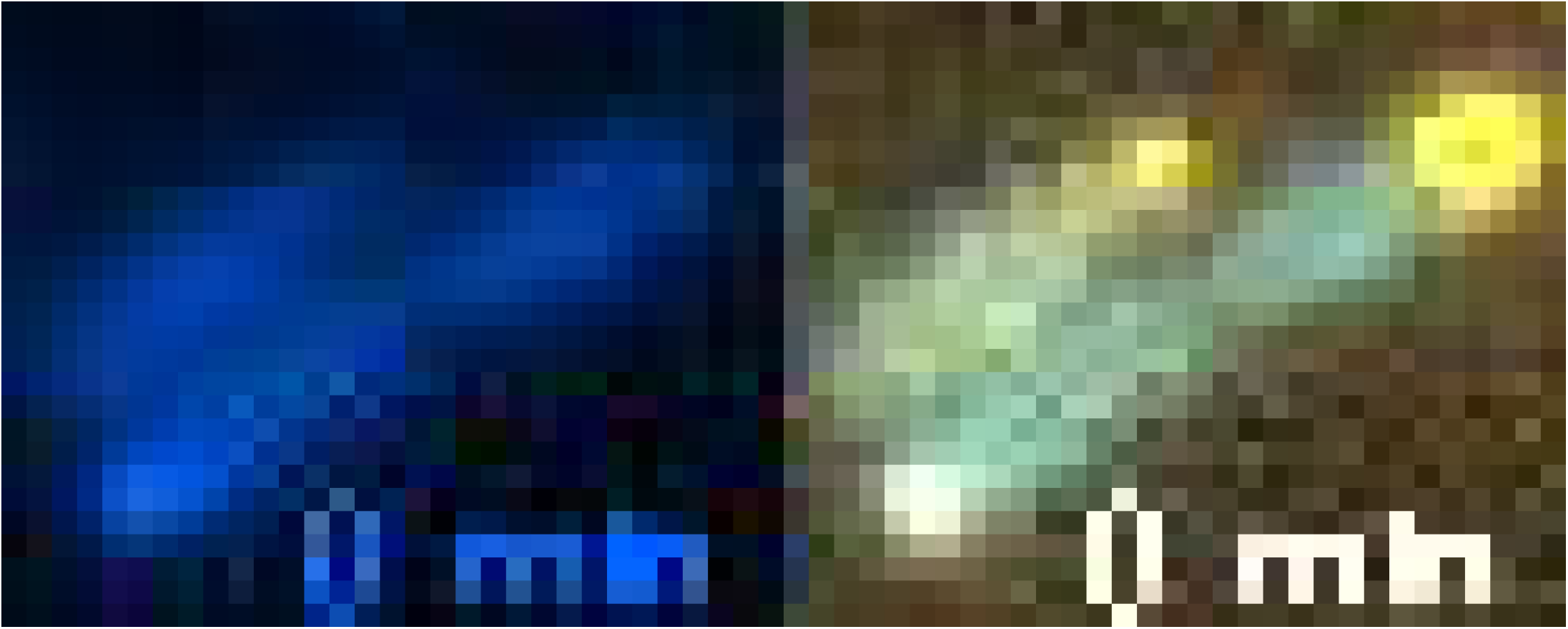

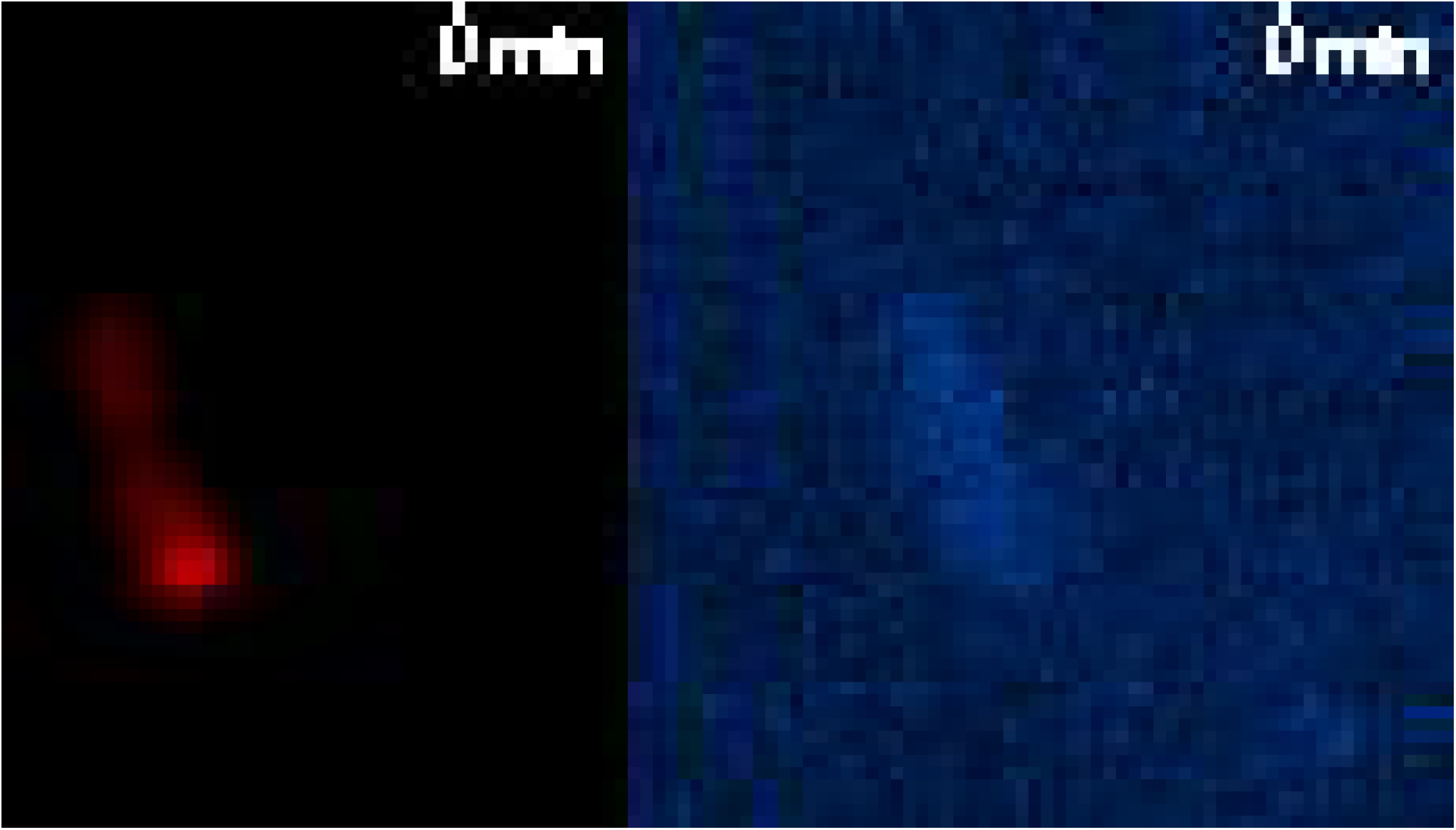

